# Single Molecule Diffusion Studies Reveal Rapid Translations Across Target Membrane Surfaces for both p40Phox PX domain and Class III PI3K VPS34 Complex 1, the Autophagy Lipid Kinase

**DOI:** 10.64898/2026.10.30.763023

**Authors:** Lucas Lizardos, Yohei Ohashi, Anne Marie McCombs, Jefferson D. Knight, Roger L. Williams, Joseph J. Falke

## Abstract

Single molecule analyses are providing new insights into the molecular mechanisms by which peripheral membrane proteins approach and translate across a target bilayer surface before stable docking. Here we compare and contrast the translational movements of four diffusers on a representative supported lipid bilayer: (i) the fluorescent lipid LRB-(DO)PE, (ii) the G-protein Rab1A anchored to (DO)PE; (iii) the PI(3)P sensor p40Phox PX domain, and (iv) the autophagy lipid kinase VPS34 Complex 1. The latter enzyme is a Class III PI-3-Kinase and its unusual architecture exhibits two distinct membrane-docking “feet” unique to Class III PI3Ks. Notably, the membrane contact surfaces of the PX domain and autophagy lipid kinase each possess multiple positive charges suitable for electrostatic interactions with their anionic target membrane. The present single molecule tracking studies reveal that both the fluorescent lipid and the lipid-anchored G protein Rab1A exhibit 2-dimensional translation typical of simple lipid diffusion in the bilayer. Single molecule density measurements indicate both PX domain and autophagy lipid kinase exhibit significantly reduced membrane docking when the bilayer negative charge is decreased, confirming the importance of their protein-bilayer electrostatic interactions. Single molecule tracking studies reveal that both PX domain and autophagy lipid kinase exhibit hopping movements across the bilayer surface during their initial bilayer contacts, yielding a significantly higher frequency of long range displacements than allowed by lipid 2-D diffusion. After transitioning from hopping to a stable bilayer-docked state, both PX domain and autophagy lipid kinase exhibit 2-D diffusion slower than lipid diffusion indicating the appearance of increased frictional drag against the bilayer due to new protein-bilayer contacts. PX domain requires its target lipid for the transition from electrostatic hopping to stable bilayer docking, while autophagy lipid kinase can dock spontaneously to anionic lipid bilayers even in the absence of its substrate lipid. In principle, the slowly diffusing, bilayer-docked states of PX domain and autophagy lipid kinase could be interrupted by unbinding-hopping-rebinding events. However, such events are not detected for PX domain in the presence of target lipid (DO)PI(3)P nor for autophagy lipid kinase in the presence of substrate lipid (DO)PI. Instead, unbinding-hopping-rebinding events are only observed for autophagy lipid kinase in the absence of substrate lipid, indicating that the presence of a specific binding lipid reduces the frequency or spatial amplitude of such events. Overall, the findings provide evidence that p40Phox PX domain possesses a search mechanism involving electrostatic hopping that speeds its acquisition of target lipid, as previously observed for PH domains. The evidence indicates that the autophagy lipid kinase also exhibits electrostatic hopping/skating behavior that is proposed to play a functional role by speeding regulatory encounters with autophagy lipid kinase activators and effectors. Moreover, the ability of the lipid kinase to spontaneously dock to anionic membranes without a target lipid could facilitate its distribution across broad regions of the growing autophagosome. Finally, the autophagy lipid kinase exhibits multiple translational modes that we hypothesize are enabled by its two membrane docking feet.

## INTRODUCTION

In the autophagy and endocytic pathways, Class III PI-3-kinases VPS34 Complex 1 and 2, respectively, phosphorylate the substrate lipid phosphatidylinositol (PI) to produce the signaling lipid phosphatidylinositol-3-phosphate (PI(3)P). The resulting PI(3)P signal drives the assembly of signaling pathways on the cytoplasmic surfaces of autophagosomes and endocytic vesicles such as phagosomes, respectively (Hurley 2026; Dessus 2026 et al; Safaroghli-Azar et al 2023). In mammalian cells, both the autophagy lipid kinase VPS34 Complex 1 and the endocytic lipid kinase VPS34 Complex 2 require a G-protein activator to stimulate substantial PI(3)P production, for example Rab1A and Rab5A, respectively (Spokaite et al 2026; Scott & Burke 2026; Tremel et al 2021; Buckles et al 2020). The autophagy lipid kinase possesses multiple activators beyond Rab1A including ULK1, GABARAP and NRBF2 (Dessus et al 2026; Chen & Hurley 2025; Mercer et al 2021; Young et al 2016; Ohashi et al 2016). Downstream components recruited to the membrane by PI(3)P possess a PI(3)P targeting domain such as PX, FYVE, PROPPIN, or PH (Rau & Haucke 2026; Montano-Rendon et al 2021).

Like other lipid signaling pathways, the autophagy pathway and endocytic pathways such as phagocytosis use the lipid bilayer as a signaling platform where peripheral proteins bind, diffuse in two dimensions, and form signaling complexes essential for pathway regulation and information transfer. The dynamics of pathway function can be influenced by translational movements on the bilayer surface that occur both during transient, initial membrane contacts and during the ensuing membrane-docked state. In principle, the effects of translational movements on pathway kinetics are increased when the density of binding partners is low. The autophagosome membrane is the fastest growing membrane in the cell with the lowest protein:lipid ratio (Melia et al 2020) and thus provides sparse protein regions where rapid translational movements of autophagy lipid kinase may speed interactions with its activator and effector proteins.

Here we investigate the 2-D movements of four representative diffusers from the autophagy and phagocytosis pathways as they translate across a membrane surface. The diffusers include a phospholipid, a phospholipid-anchored G protein, a PI(3)P-binding PX domain, and an active, lipid-associated Class III PI3K lipid kinase enzyme (**Figure 1A**). (i) LRB-PE is a fluorescent phospholipid possessing Lissamine rhodamine B (LRB) covalently coupled to the headgroup of PE, a standard background lipid in autophagosome and phagosome membranes. (ii) The lipid-anchored G protein Rab1A is covalently coupled to the headgroup of a maleimide-PE lipid via its C-terminal Cys residue that is natively lipidated in cells. (iii) The p40Phox PX domain functions to recruit its parent protein to PI(3)P on the phagosome membrane where it activates NADPH oxidase complex to generate an ROS burst that inactivates trapped pathogens; here we employ the isolated PX domain as a PI(3)P sensor (Suh et al 2006). (iv) The autophagy lipid kinase VPS34 Complex 1 is a heterotetrameric Class III PI-3-kinase formed by the association of four subunits: VPS34, VPS15, Beclin1, and ATG14L. This lipid kinase binds reversibly to the target bilayer and generates PI(3)P with a modest intrinsic activity until boosted by association with membrane-anchored GTP-Rab1A and/or with another activator (Hurley 2026; Dessus et al 2026; Tremel et al 2021; Ohashi 2021a,b). The resulting PI(3)P signal initiates and drives the growth of the autophagosome double membrane and may activate other signaling pathways.

**Figure 1.**
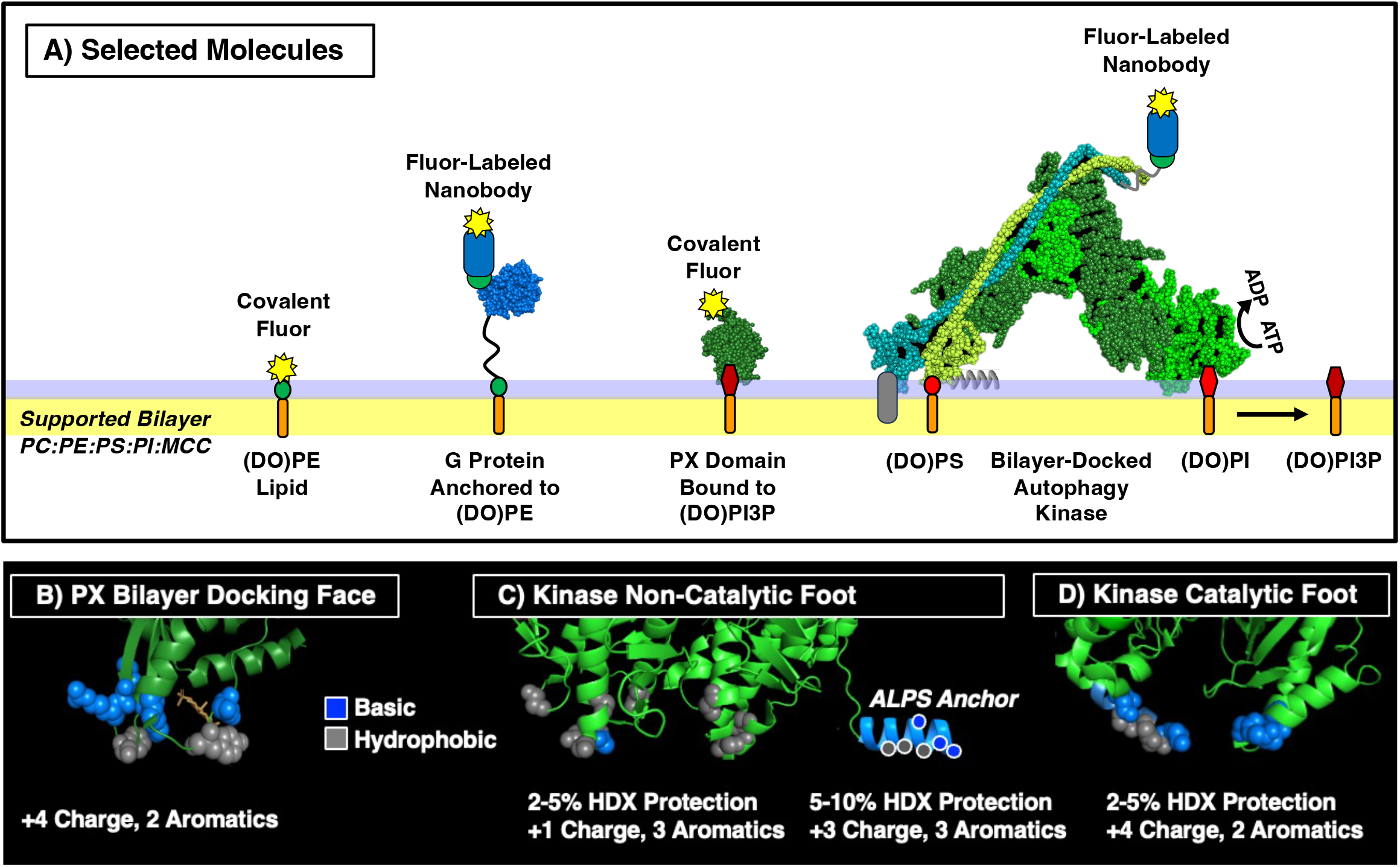
Membrane-bound molecules utilized in this study, and key protein-bilayer contact surfaces. **A)** Shown are the four fluorescent molecules studied: (i) phospholipid (DO)PE covalently labeled with fluor on its headgroup, (ii) G-protein Rab1a (Zhang et al 2024) covalently anchored to the bilayer via coupling of its native lipidation Cys to a (DO)PE headgroup, then fluor-labeled on demand by addition of fluorescent nanobody, (iii) p40Phox PX domain (Bravo et al 2001) covalently labeled with fluor and bound to its target lipid (DO)PI(3)P, and (iv) Class III PI-3-Kinase VPS34 Complex 1 (Cook et al 2025) docked to a target bilayer then fluor-labeled when desired by addition of fluorescent nanobody (Gotzke et al 2019). While the models presented herein show autophagy lipid kinase “standing” on the membrane with the molecular plane perpendicular to the bilayer (Ma et al 2017), it is likely this docking angle fluctuates. **B-C)** Shown are the predicted, bilayer-docked configurations of **B)** PX domain with its membrane docking loops, **C)** the non-catalytic foot of autophagy lipid kinase highlighting its proposed membrane docking motifs (aromatic finger, bound anionic lipid, and the essential ALPS helix), and **D)** the catalytic foot of autophagy lipid kinase showing the substrate cleft bordered by the essential C-terminal basic helix and other basic residues. Basic residues are blue and aromatic residues are gray; net positive charge is estimated. For autophagy lipid kinase, also shown are bilayer-induced, partial protections from hydrogen-deuterium exchange measured for the indicated regions, providing evidence that both feet contact the bilayer (Ohashi et al 2020).

The PX domain and the autophagy lipid kinase are both peripheral membrane docking proteins that transition from a cytoplasmic inactive state to an active, membrane-bound state. Thus, both have membrane docking surfaces that drive their protein-membrane interactions. **Figure 1B** shows the membrane-docking face of the PX domain (Bravo et al 2001), which is computationally modeled to contact the membrane via two loops while PI(3)P occupies the headgroup binding cleft (Lomize et al 2012). The modelled interaction surface possesses a net positive charge of as well as 2 aromatic sidechains that may intercalate into the bilayer.

The remarkable two-legged structure of the autophagy lipid kinase, unique to Class III PI-3-kinase enzymes, possesses a membrane docking face on each of its two “feet” (Cook et al 2025; Ohashi et al 2020). These membrane docking surfaces display positive charges and aromatic residues believed to stabilize the protein-bilayer complex. Hydrogen-deuterium exchange studies of the autophagy lipid kinase have shown that the non-catalytic foot and the catalytic foot both contact the bilayer in the membrane-docked state, yielding partial protection from aqueous exposure (Ohashi et al 2020). The non-catalytic foot (**Figure 1C**) possesses a net positive charge, much of which is located on the C-terminal, amphiphilic ALPS helix of the ATG14L regulatory subunit (Fan et al 2011). This foot also possesses multiple hydrophobic residues on the ALPS helix and on the aromatic finger of the Beclin 1 regulatory subunit. Similarly, the catalytic foot (**Figure 1D**) possesses a net positive charge, much of which resides on the C-terminal helix of the VPS34 catalytic subunit, as well as several hydrophobic residues. Both the ALPS helix of the non-catalytic foot and the C-terminal helix of the catalytic foot are known to be absolutely essential for lipid kinase activity (Ohashi et al 2020; Brier et al 2019; Stjepanovic et al 2017). Models presented herein show autophagy lipid kinase “standing” on the membrane with the molecular plane perpendicular to the bilayer (Ma et al 2017), although fluctuations in this docking angle are likely.

Because the membrane docking faces of the PX domain and the autophagy lipid kinase are each positively charged, we hypothesize that both proteins are capable of an electrostatic search of the membrane surface prior to stable docking to the target bilayer, as previously observed for PH domains (Corbin et al 2004; Knight & Falke 2009; Lai et al 2013). **Figure 2** schematically illustrates an electrostatic search culminating in binding to a target, in this case a lipid. During the initial contacts with the anionic membrane surface, the positively charged membrane docking face of the experiences electrostatic attraction to the anionic membrane enabling the protein to stochastically bounce or hop along the surface until it encounters its binding target. During the hopping mode the protein moves primarily through the aqueous phase with only transient bilayer contacts, thus it can translate across the bilayer surface faster than lipids diffuse in the bilayer where the viscosity is 2-3 orders greater (Adrian et al 2022, Ziemba & Falke 2013). After stable bilayer docking is achieved, frictional drag against the bilayer dominates and the protein diffuses at the speed of a lipid, or slower than lipid if the protein itself forms bilayer contacts that increase frictional drag. Rigorous studies have shown that the diffusion constants of proteins docked to supported lipid bilayers of the same lipid composition are inversely proportional to the total frictional drag of the protein-lipid complex against the bilayer (Knight et al 2010; Ziemba et al 2012; Ziemba & Falke 2013; Camley et al 2015).

**Figure 2.**
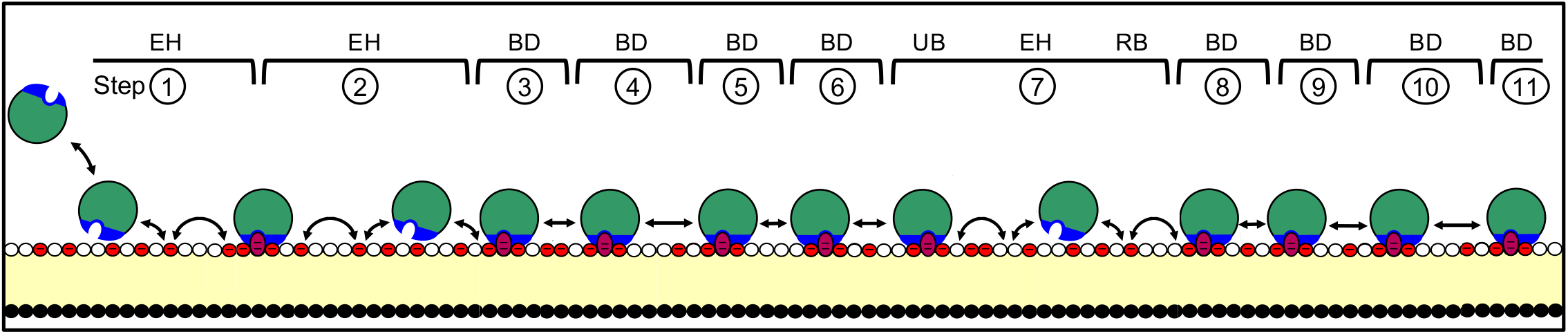
Schematic single molecule trajectory of a fluorescent peripheral membrane docking protein encountering an anionic bilayer and binding to its target lipid. The illustration shows a single molecule trajectory, linearized for simplicity, as captured in a series of numbered, consecutive 20 msec video frames. Schematic, microscopic protein translational movements occur during each frame. The membrane docking face of the protein is positively charged (blue) while the bilayer is negatively charged due to the presence of anionic lipid headgroups, predominantly those of phosphatidylserine (red circles). During the initial bilayer contacts the protein is proposed to translate via faster-than-lipid, long-range electrostatic hops (EH) across the bilayer surface, speeding the on-rate for binding to its target lipid. After target lipid docking, the bound protein diffuses (BD) in 2-dimensions in the plane of the bilayer. It may undergo an unbinding-hopping-event (UB-EH-RB) that either yields dissociation (UB-EH-D) or rebinding to the same or different target lipid molecule (UB-EH-RB), thereby extending the trajectory. Such an electrostatic search mechanism has been documented for PH domains (Corbin et al 2004; Knight & Falke 2009; Lai et al 2013) and the present findings support similar electrostatic search mechanism for PX domain and autophagy lipid kinase.

Single molecule TIRFM studies of reconstituted signaling reactions on supported lipid bilayers can provide unique insights into membrane-based signaling pathways (Duewell 2026; Groves 2025; McCombs et al 2024; Gordon et al 2023; Buckles et al 2017; Ziemba et al 2016; Knight & Falke 2009; Wagner & Tamm 2001). The present study employs existing and new single molecule TIRFM methods to directly observe and quantify the movements of each diffuser in its bilayer-associated state. Each single molecule diffusion trajectory is obtained by imaging a fluorescent diffuser in a series of consecutive 20 msec video frames (Figure 2). Since the molecule moves during the exposure, often yielding a diffuse molecular “spot”, the particle tracking software identifies the location of the molecule in that frame as the brightness-weighted intensity centroid (Sbalzarini & Koumoutsakos 2005; Ershov et al 2022). The displacement between the centroid in consecutive frames defines the length of a “step” between them. Note that each 20 ms “step” is actually a 20 ms “walk” during which the molecule executes a vast number of stochastic, Brownian micro-steps on the sub-molecular length scale, together yielding the net displacement “step” between consecutive frames. While lipid-anchored diffusers are coupled to the bilayer throughout their trajectories, a membrane docking protein trajectory begins with a target membrane encounter yielding transient protein-membrane contacts, which may include electrostatic hopping during an electrostatic search of the bilayer surface to acquire a target lipid or protein. Subsequently, the protein undergoes stable docking to the bilayer and begins lipid-associated 2-D diffusion that may be disrupted by an unbinding-hopping-rebinding event. Finally, the molecule disappears due to full dissociation (or, less frequently, due to bleaching or diffusion out of the field).

Overall, the present findings reveal that both the p40Phox PX domain and the autophagy lipid kinase can rapidly translate across the surface bilayer faster than possible for the lipids diffusing within the bilayer. Such rapid translation may well increase the efficiency of their signaling on phagosome and autophagosome membranes, respectively. For both proteins, the fast translational components peak early in their trajectories soon after the initial membrane contact, then decay as the trajectories relax into their bilayer-docked mode. In their bilayer-docked state both proteins exhibit 2-D diffusion slower than lipids in the same bilayer, indicating the appearance of new protein-bilayer contacts that increase the frictional drag of the membrane-docked protein against the bilayer. When no substrate lipid is available, the membrane-docked autophagy lipid kinase occasionally exhibits unbinding-hopping/skating-rebinding events, yielding detectable long-range displacements in trajectories (Figure 2).

## MATERIALS AND METHODS

### Reagents

Synthetic dioleolyl phospholipids PC (phosphatidylcholine; 1,2-dioleoyl-sn-glycero-3-phosphocholine, SKU 850375C, CAS 4235-95-4), PS (phosphatidylserine; 1,2-dioleoyl-sn-glycero-3-phospho-L-serine, SKU 840035C, CAS 90693-88-2), PE (1,2-dioleoyl-sn-glycero-3-phosphoethanolamine, SKU 850725C, CAS 4004-05-1), PI (1,2-dioleoyl-sn-glycero-3-phosphoinositol, SKU 850149P, CAS 799268-53-4), PI(3)P (1,2-dioleoyl-sn-glycero-3-phospho-(1’-myo-inositol-3’-phosphate), SKU 850150P, CAS 1246303-09-2), LRB (DOPE-Liss Rhod, CAS 384833-00-5) and the maleimide-modified headgroup phospholipid PE-Mal (1,2-dioleolyl-sn-glycero-3-phosphoethanolamine-N-[4-(p-maleimidomethyl)cyclohexane-carboxamide], SKU 780201, CAS DO-0365253) were from Avanti Polar Lipids (Alabaster, AL). Alexa Fluor 555 C2 maleimide (A20346, Thermo Fisher Scientific, Waltham, MA). Sulfuric acid (ACS Grade, 95.0–98.0% purity, H_2_O_4_S, CAS 7664-93-9) and sodium hydroxide (Certified ACS pellets, ≥97.0% purity, HNaO, CAS 1310-73-2) were bought from Fisher Chemical (Pittsburgh, PA). Hydrogen peroxide (30% solution, AR grade for ACS analysis, H_2_O_2_, CAS 7722-84-1) was obtained from Avantor Macron (Radnor, PA). Potassium hydroxide (≥85.0% purity, HKO, CAS 1310-58-3) was sourced from Pittsburgh, PA. Adenosine 5’-triphosphate disodium salt hydrate (ATP-Na2, C_10_H_14_N_5_Na_2_O_13_P_3_ ⋅ xH_2_O, CAS 34369-07-8), magnesium chloride hexahydrate (99.995% trace metals basis, Cl_2_Mg ⋅ 6H_2_O, CAS 7791-18-6), sodium chloride (Halite, 99.999% trace metals basis, ClNa, CAS 7647-14-5), potassium chloride (99.999% trace metals basis, ClK, CAS 7447-40-7), and HEPES (≥99.5% titration, Cat. No. H3375) were obtained from Sigma-Aldrich (St. Louis, MO). Reduced L-glutathione (≥98.0% purity, Cat. No. G4251-10G), bovine serum albumin (BSA, CAS 9048-46-8, Cat. No. A7030-10G), and manganese(II) chloride tetrahydrate (Cat. No. 203734-5G) were sourced from Sigma-Aldrich Fine Chemicals Biosciences (St. Louis, MO).

### Materials

Milli-Q water filtered with a Quantum TEX filter (part no. QTUM0TEX1) was obtained from MilliporeSigma (Guyancourt, France). Hourglass HandPRO Fortis500 nitrile gloves with advanced cross-linking (X-linked, ref. 503) were sourced from Hourglass International (Denver, CO, USA). Black diamond weighing boats (5 mL, 35 x 55 mm, part no. J6501B) were obtained from Alkali Scientific (Fort Lauderdale, FL, USA). Sartorius Vivaspin 500 centrifugal concentrators (10 kDa MWCO, PES ultrafiltration membrane, 0.5 mL; part no. 14558396) were obtained from Sartorius (Göttingen, Germany). TipOne 1000 µL aerosol barrier filter pipette tips were acquired from USA Scientific, Inc. (Ocala, FL, USA). Perform 10 µL XL (SKU: PFM-10XL-FT) and 200 µL XL (SKU: PFM-200XL-FT) ultra-low retention filtered pipette tips were obtained from Life Science Products (Chestertown, MD, USA). Cover-Well perfusion chamber gaskets (catalog no. C18139) were purchased from Invitrogen (Carlsbad, CA, USA). Borosilicate micro cover glasses (22 × 22 mm × 0.16–0.19 mm thick, non-sterile, part no. NC1466408) were obtained from Ted Pella Inc. (Redding, CA, USA). Axygen 2 mL self-standing conical microcentrifuge tubes with brown screw caps and O-rings (sterile, polypropylene, RNase/DNase- and pyrogen-free, rated to 20,000 × g RCF) were acquired from Corning (Union City, CA, USA). Thermo Scientific 15 mL (part no. 339650) and 50 mL (part no. 339652) conical centrifuge tubes (bulk, screw cap, graduated, single-use, RNase/DNase-free, max RCF 17,000 × g) were acquired from Thermo Fisher Scientific (Pittsburgh, PA, USA). Additionally, glass Pasteur pipettes (5.75 in., part no. 77574-984), Fisherbrand 0.5 mL microcentrifuge tubes (locking snap cap, polypropylene, DNase/RNase- and pyrogen-free, non-sterile, autoclavable, rated to −86 °C and 30,000 × g RCF), and mdi PVDF syringe filters (13 mm, 0.2 µm, female luer lock/male luer slip, HPLC certified, non-sterile, catalog no. SYVF0301MNXX104) were sourced from Fisher Scientific (Pittsburgh, PA, USA).

### Protein Preparation

ALFA-tagged autophagy lipid kinase (ALFA-VPS34 C1) was expressed and purified as previously described for untagged VPS34 C1 (Ohashi et al. 2016; Ohashi et al 2020; Tremel et al 2021) with the following modifications. A plasmid carrying an ALFA tag coding fragment at the N-terminus of *BECN 1* coding region was constructed (pYO1524) and co-transfected with pYO1025 (Ohashi et al, 2016 Addgene #235705) and pYO1100 (Tremel et al 2021, Addgene #235707) into 3.2 L Expi293f suspension cells (ThermoFisher A14527) and were grown in Expi293 Expression Medium (ThermoFisher A1435102) at 37°C, 8% CO_2_, shaking at 125rpm. At density of 2×10^6^/L, cells were transfected with 1.1 mg DNA/L cells using polyethyleneimine ‘MAX’ (Polysciences 24765, 1mg/mL in PBS) at 3mg/L. Cells were grown further in the same conditions as above for 68 hours, then harvested by centrifugation at 3000 g for 15 min, flash frozen in liquid nitrogen, and stored at −80 °C until use. To purify the protein, cells were resuspended in 50 mL/2 L cells of lysis buffer (50 mM HEPES pH 8.0, 150 mM NaCl, 1% Triton X-100 (Fisher BioReagents BP151), 12% glycerol, 2 mM MgCl₂, 1 mM phenylmethylsulfonyl fluoride (PMSF, Thermo Fisher Scientific 36978), 0.5 mM tris(2-carboxyethyl)phosphine (TCEP), and 1x EDTA-free protease inhibitor cocktail tablet (Roche 11873580001)). Lysate was cleared by centrifugation at 142,000 g for 30 min at 4 °C in a Type 45 Ti rotor (Beckman Coulter) and filtered through a 5 µm syringe filter (Sartorius 17594). The filtered lysate was incubated for 3.5 hours at 4 °C and 8 rpm rotation with 2 mL IgG beads (Cytiva 17-0969) equilibrated with lysis buffer. Beads were then transferred to a gravity column, washed with 50 mL ice-cold wash buffer (50 mM HEPES pH 8.0, 150 mM NaCl, 0.1% Triton X-100, 12% glycerol, 50 mM MgCl₂, 0.5 mM TCEP, 4 µg/mL bovine pancreatic RNase (Sigma 83834)), and incubated with 5 mL wash buffer supplemented with 5 mM ATP for 20 minutes. Beads were further washed with another 50 mL wash buffer and 100 mL TEV buffer (50 mM HEPES pH 8.0, 150 mM NaCl, 0.5 mM TCEP), then resuspended in 10 mL TEV buffer and incubated with 40 µL of 4.4 mg/mL TEV protease (house made) overnight in the cold room. Flow-through was collected and remaining protein was eluted three times with 5 mL TEV buffer followed by 7 mL TEV buffer supplemented with 0.1% CHAPS (Calbiochem 220201). Eluted fractions were combined, passed through a 0.22 µm syringe filter (Elkay E25-PS22-50S), concentrated to 0.5 mL using a 100k MWCO concentrator (Millipore UFC810024), and injected onto an S200 10/300 GL size exclusion chromatography column equilibrated with 20 mM HEPES pH 8.0, 150 mM NaCl, 0.5 mM TCEP. The main peak fractions were pooled, concentrated, frozen in liquid nitrogen in 2 µL aliquots, and stored at −80°C until use.

ALFA-Rab1A was expressed and purified as previously described for untagged Rab1A (Tremel et al 2021) with the following modifications. The ALFA tag coding fragment was introduced at the N-terminus of RAB1A Q70L C26S C126S coding region (Tremel et al 2021). The resulting plasmid (pYO1726) was transformed into *E.coli* OVEREXPRESS C41(DE3) chemically competent cells (Lucigen 60442). Transformed bacteria were cultured in 4 L 2xTY medium supplemented with 0.1 mg/mL ampicillin (Formedium AMP50) at 37 °C until OD_600_=0.3, then expression was induced with 0.5 mM IPTG, and bacteria were incubated for further 19 hours at 16 °C. Cells were pelleted by centrifugation at 6700 g for 15 minutes and flash frozen in liquid nitrogen, stored at −80 °C until use. The pellets from 2 L culture were resuspended in 50 mL lysis buffer (25 mM HEPES pH 8.0, 200 mM NaCl, 12% glycerol, 5 mM DTT, 1 mM PMSF, 1x EDTA-free protease inhibitor cocktail tablet). Cells were sonicated for 5 min on ice (2 s on, 3 s off, 60% amplitude), and lysate cleared by centrifugation at 142,000 g for 15 min at 4 °C in a Type 45 Ti rotor. Supernatant was filtered through a 5 µm syringe filter and incubated with 2.5 mL GS4B-GST beads (Cytiva 17-0756) for 2 hours at 4 °C at 8 rpm rotation. Beads were then transferred to a gravity flow column, washed with 100 mL lysis buffer, 100 mL wash buffer (25 mM HEPES pH 8.0, 300 mM NaCl, 20% glycerol, 5 mM EDTA, 5 mM DTT), and 50 mL TEV buffer (25 mM HEPES pH 8.0, 150 mM NaCl, 5 mM DTT). Beads were resuspended in 10 mL TEV buffer, then 70µL of TEV protease at 3 mg/mL was added, and sample were incubated overnight without rotation to cleave the N-terminal GST-tag. Flow-through was collected and further eluted with 20 mL HepA buffer (25 mM HEPES pH8.0, 1 mM TCEP, 20 mM imidazole, 1 mM PMSF, 1x EDTA-free protease inhibitor cocktail tablet). The elution fractions were combined, and the total volume was adjusted to 50 mL with HepA buffer. The protein solution was passed through a 0.22 µm syringe filter, then passed through a 5 mL HiTrap heparin column (Merck, GE17-0406-01) to remove TEV protease. The flowthrough fraction was concentrated down to 900 µL at 10 mg/mL (449 µM) using a 10 k cut off concentrator (Millipore, UFC901024). Concentrated protein was incubated with 11 molar excess GTP or GDT and 18 molar excess EDTA for 90 min at room temperature, then MgCl_2_ was added at the final concentration of 20 mM and further incubated for 30 min at room temperature. The incubated samples were loaded onto a Superdex75 16/60 column that had been equilibrated with GF buffer (25 mM HEPES 8.0, 150 mM NaCl, and 0.5 mM TCEP). Peak fractions were collected and concentrated using 10k cut-off concentrator.

p40Phox PX domain was expressed and purified previously described (Ohashi et al, 2020). Subsequently PX domain was labeled at an N-terminal native Cys residue (Cys 5) with Alexa Fluor 555 as follows. Protein was diluted to 100 µM in 450 µL of labeling buffer (20 mM HEPES pH 7.2 with KOH, 150 mM KCl, 0.5 mM fresh TCEP). Avoiding fluor exposure to bright light, 1 mg probe was dissolved in 200 µL anhydrous yielding a 4 mM probe stock. To the protein sample, probe stock was added to a final concentration of 100 µM and gently pump mixed with a pipettor. The labeling reaction was wrapped in foil and incubated with slow rotation at 72° C for 20 min. Unreacted probe was quenched by adding glutathione to 1 mM with gentle pump mixing, re-wrapping in foil and incubating with slow rotation at 72° C for 15 min. The labeled protein was snap frozen in 50 µL aliquots and stored at −80° until washing the protein to remove unbound probe. After thawing on ice, 100 µL protein was diluted with labeling buffer to 500 µL in a pre-rinsed 10,000 MW cutoff Sartorius Vivaspin 500 spin concentrator. The concentrator was spun at 10,100 x g at 4° C for 8 min down to 100 µL, discarding the flow through then repeating 4x. The resulting 100 µL of washed protein was subjected to nanodrop concentration analysis (A280, A555) and diluted to 1000 µL with labeling buffer before aliquoting and storage as below.

Fluorescent anti-ALFA nanobody labeled with 2 molecules of AZ568 fluor was obtained commercially as a lyophilized powder (FluoTag-X2 anti-ALFA single domain nanobody, N1502-AF568-L, Abnova, Taoyuan City, Taiwan). The powder was reconstituted in glycerol:H2O 1:1 (v:v) as per Abnova instructions. The equilibrium dissociation constant for anti-ALFA nanobody binding to ALFA-tagged protein was measured to be 6 µM.

Each protein was aliquoted in a specific storage buffer, snap frozen in liquid N2, and stored at −80° C as follows before thawing on ice just before use. ALFA-tagged autophagy lipid kinase (2.7 µM) in 20 mM HEPES 7.9, 150 mM NaCl, 0.5 mM TCEP was snap frozen in 6 µL aliquots. ALFA-tagged GTP-Rab1A (5.3 µM) and GDP-Rab1A (4.7 µM) in 20 mM HEPES 7.0, 150 mM NaCl, 5 mM TCEP, 2 mM MgCl2 were snap frozen in 4 µL aliquots. AF-555 labeled PX domain (10 µM) in 20 mM HEPES pH 7.2 with KOH, 150 mM KCl, 0.5 mM fresh TCEP was snap frozen in 10 µL aliquots. FluoTag-X2 anti-ALFA nanobody in glycerol:H2O 1:1 (v:v) was snap frozen at 2.5 µM in 5 µL aliquots.

### Supported Lipid Bilayer Preparation

All steps preparing samples for single molecule TIRFM employed ultrapure reagents and materials to avoid fluorescent contaminants, which are found at unacceptably high levels in many standard laboratory supplies.

Homogeneous lipid mixtures were prepared using (DO)PC/(DO)PE/(DO)PS/(DO)PI/DOMCC at a 50:18:25:5:2 molar ratio. First, the individual lipids dissolved in chloroform were mixed to yield the desired composition and molar ratio and aliquoted into 2 mL amber screw-top tubes. The solvent was removed by SpeedVac without heating until the pressure was below 180 mTorr. The resulting lipid film was rehydrated to a total lipid concentration of 2-3 mM with 0.2 µm-filtered buffer containing 25 mM HEPES, 140 mM KCl, 15 mM NaCl, 0.5 mM MgCl_2_, and 2 mM TCEP, pH to 7.2 with KOH. Resuspension of the film included pump mixing and scraping the film from the walls of all tubes with the tip of a 1 mL pipettor, followed by vortexing each tube for 1 min and snap freezing them together in liquid N2. All tubes were thawed in a rotating 40° C water bath, then resuspension was completed by again pump mixing, scraping and vortexing each tube for 1 min. The tubes were combined 0.7 mL aliquots of the rehydrated lipid mixture in 2 mL amber screw-top tubes were snap-frozen in liquid N_2_ and stored at −80 °C until use.

Where indicated in the figure legends, the above lipid mixture was modified to remove or add specific lipids in the chloroform mixture, or to add covalently coupled Rab1A. When (DO)PS or (DO)PI was removed from the lipid mixture it was replaced with equivalent moles of (DO)PC. Experiments with fluorescent lipid included 8 ppm (DO)PE-LRB. Experiments with PX domain included 0.6% −0.8% mole percent (DO)PI(3)P, except in autophagy kinase experiments wherein the kinase generated product (DO)PI(3)P. Experiments with Rab1A included an overnight lipid coupling step (see below).

Glass coverslips were ultra-cleaned by treatment with piranha solution, a 3:1 ratio of concentrated sulfuric acid to hydrogen peroxide, by placing 9 drops of concentrated sulfuric acid in the center of each cover slip using a clean Pasteur pipette, adding 3 more drops of hydrogen peroxide with a new pipette, followed by a 1 hr incubation at 23.0 ± 0.5 °C. Noting which side was treated, each coverslip was then rinsed 6+ times on each side with 15 total mL of Milli-Q water, dried by blowing compressed N_2_ gas, and exposed to an ozone chamber for 9 h.

On the day of the experiment, a sheet of Coverwell rubber gaskets was cut into individual gaskets. Then each was rinsed three times with Milli-Q water, blotted with a Kimwipe and air dried for 5 min. Subsequently each was pressed onto an ultra-cleaned glass coverslips to ensure a secure seal yielding a ∼50 mL chamber with two entry-exit ports.

The same day, each desired lipid aliquot was thawed on ice, vortexed for one minute, suspended in an ice water bath and sonicated using a Qsonica Q700 sonicator equipped with a microtip 3.2 mm (part No. 4418) for a total of 4 min (Pulse-ON time: 0.1 s, Pulse-OFF time: 0.2 s). The resulting small unilamellar vesicles (SUVs) were mixed 1:1 with a 0.2 µm-filtered deposition buffer containing 25 mM HEPES, 140 mM KCl, 0.5 mM MgCl_2_, and 0.5 M NaCl, pH to 7.2 with KOH. This 1:1 mixture was deposited on an ultra-clean glass surface inside the imaging gaskets and incubated for 45 min.

Following deposition, the lipids were rinsed with approximately 15 mL total Milli-Q water, alternating between the two ports, and taking care to avoid introducing bubbles, which can destroy the bilayer, into the chamber. Each imaging gasket was then rinsed by gentle pump mixing with a 50 µL aliquot of coupling buffer containing containing 25 mM HEPES, 150 mM NaCl, pH to 7.4 with NaOH then repeating 4x. Experiments with lipid-anchored Rab1A, included a lipid coupling step as follows: for each gasket, most of the residual liquid was removed from the ports, then 40 µL of coupling buffer containing 5 µM Rab1A was injected and the solution was pump-mixed 5 times with the existing 40 µL of buffer, yielding an approximate 2.5 µM final concentration of Rab1A in the gasket. The resulting gaskets were incubated in a tightly sealed Pyrex glass humidity chamber containing Milli-Q water-moistened Kimwipes to keep the gaskets from drying out at 4 °C overnight for 16-20 hours for experiments involving Rab1A, or for 90 min at 23.0 ± 0.5 °C for experiments employing the other diffusers. Next each imaging gasket was rinsed by gentle pump mixing with a 50 µL aliquot of pH 7.4 imaging/activity buffer containing 25 mM HEPES, 75 mM KCl, 75 mM NaCl, 2.0 mM Mg^2+^, 75 µM Mn^2+^, and 5 mM glutathione then repeating 4x, followed by incubation at 23.0 ± 0.5 °C for 60 min. No BSA was present at this step to prevent the coupling of its sulfhydryl or amino groups to the maleimide. The gaskets were then rinsed 5 more times with the same buffer supplemented with 10 µg/mL BSA to block bilayer defects that could otherwise yield localized patches that bind proteins nonspecifically. The gaskets were then ready to use for smTIRFM.

### Preparing (DO)PE-LRB Samples for Imaging

Imaging/activity buffer (+) BSA (+) ATP was prepared containing final concentrations 10 µg/mL BSA,1mM Mg-ATP, pH 7.4 with gentle mixing by inversion to avoid foaming the protein. Imaging/activity buffer (+) BSA (+) ATP was then used to wash the gaskets prior to imaging as follows. Using a pipettor set to 42.5 µL, the residual liquid sitting on the surface of the gasket above the ports was removed. Next, 42.5 µL of imaging/activity buffer (+) BSA (+) ATP was injected into each gasket without pump mixing taking care to avoid the introduction of air bubbles, and the excess volume extruded from the ports was removed. Finally, an additional 42.5 µL of the imaging/activity buffer was injected into each gasket without pump mixing then smTIRFM imaging of the gasket was carried out.

### Preparing AF555-PX Domain Samples for Imaging

To prepare an AF555-PX domain stock solution, first a protein dilution buffer was made by adjusting the above coupling buffer to pH 8.0 with addition of BSA and glutathione to 10 µg/mL and 1 mM, respectively. A 10 µL aliquot of 10 µM PX domain stock was thawed and serially diluted with ice cold protein dilution buffer, with all mixing provided by gentle pump mixing with a pipettor, to achieve a final 16 µM PX domain stock that was stored on ice. Just prior to smTIRFM imaging of a set of PX domain gaskets the PX stock was diluted to 0.3-0.5 nM final concentration in room temperature (21.4 °C) imaging/activity buffer (+) BSA (+) ATP and this imaging solution was used in all gaskets.

Next each gasket was prepared individually for PX domain imaging as follows. Using a pipettor set to 42.5 µL, the residual liquid sitting on the surface of the gasket above the ports was removed. Subsequently, 42.5 µL of the 0.3-0.5 nM PX imaging solution was injected into the chamber without pump mixing, taking care to avoid the introduction of air bubbles, and the resulting excess volume extruded from the ports was removed. Then a second 42.5 µL of 0.3-0.5 nM PX imaging solution was injected with no pump mixing and smTIRFM imaging of the gasket was completed.

### Preparing AZ568-Anti-ALFA-Nanobody for Imaging Rab1A or Autophagy Lipid Kinase

To prepare the nanobody for imaging ALFA-tagged Rab1A or autophagy lipid kinase, an aliquot of 2.5 µM nanobody was thawed on ice and diluted to 90 nM final concentration with 137 mM NaCl, 2.7 mM KCl, 10 mM Na2HPO4, 1.7 mM KH2PO4, pH 7.4 then stored on ice. Just before smTIRFM imaging anti-ALFA Rab1A or autophagy lipid kinase, the 90 nM stock was diluted to 1.5 nM final concentration in room temperature (21.4 °C) imaging/activity buffer (+) BSA (+) ATP. The resulting imaging solution was stored in the dark and used in all gaskets.

### Preparing ALFA-Tagged Rab1A Samples for Imaging

To image lipid-anchored Rab1A, each gasket was prepared as follows. Using a pipettor set to 42.5 µL, the residual liquid sitting on the surface of the gasket above the ports was removed. Subsequently, 42.5 µL of the 1.5 nM nanobody imaging solution was injected into the chamber without pump mixing, taking care to avoid the introduction of air bubbles, and the resulting excess volume extruded from the ports was removed. Next a second 42.5 µL aliquot of the 1.5 nM nanobody imaging solution was injected with no pump mixing and smTIRFM imaging of the gasket was carried out.

### Preparing ALFA-Tagged Autophagy Lipid Kinase Samples for Imaging

To prepare the kinase for imaging, immediately before imaging a set of kinase gaskets a single 10 µL aliquot of 3.8 µM ALFA-tagged autophagy lipid kinase was thawed and diluted a final concentration of 153 nM with the previously described ice cold protein dilution buffer. The solution was gently pump mixed and returned to ice. Next, for a given gasket, a pipettor set to 42.5 µL was used to remove the residual liquid sitting on the surface of the gasket above the ports. Subsequently, 42.5 µL of 1.5 nM nanobody imaging solution was injected into the chamber without pump mixing, taking care to avoid the introduction of air bubbles. Next, to a tube containing 47 µL of the 1.5 nM nanobody imaging solution, the autophagy lipid kinase stock was added to 1.5 nM final kinase concentration with gentle pump mixing and 42.5 µL was injected into the gasket without pump mixing. Finally, smTIRFM imaging of the gasket was carried out.

### Single-molecule TIRFM measurements

TIRFM experiments were performed at 20.5 ± 0.5 °C using an objective based setup. The instrument featured a Nikon TE2000U inverted TIRF microscope (Melville, NY) equipped with a Nikon Apochromat 60x, NA 1.49 oil immersion TIRF objective. Excitation was provided by a 532 nm diode-pumped solid-state laser (CNI-Laser, model MGLIII-532-300 mW, CNI, Changchun, China), with power attenuated as needed by density filters (ND) totaling an absorbance of 0.0 O.D. for (DO)PE-LRB measurements, O.D. 0.6 for PX domain measurements, and O.D 1.1 for AZ568-Anti-ALFA-Nanobody measurements. Fluorescence emission was passed through a 550 nm long pass filter and captured using an Andor iXon Life electron-multiplying charge-coupled device (EMCCD) camera (Oxford Instruments, Abingdon, UK). Immediately after the final pump-mixing step, each sample-loaded gasket was transferred to the microscope stage. TIRF acquisition and focusing was performed, and the initial movie was recorded within 10 to 30 s of the final mixing. Subsequent movies were acquired by moving the stage 3-5 fields away from the last field to avoid local heating by the laser irradiation, then starting another video. This process was repeated until 6+ movies were collected for this gasket. The gasket was removed and the next gasket prepared for imaging, taking ∼1.5 min to reach the final pump mixing step before transferring to the microscope and collecting the next 6+ videos. Each 5 s video was recorded with a 20 ms exposure time for consecutive frames.

### Data analysis

For each diffuser, 3,000 - 9,000 total trajectories were combined after filtering to remove minor populations of contaminating diffusers (fast, dim trajectories) and damaged diffusers (bright, immobile spots). **Figure 3** summarizes the data analysis workflow. First, each Nikon .nd2 video file was processed in ImageJ/Fiji (version 2.16.0/1.54p) by adjusting contrast with “auto” mode and converting to .avi file format. Next, single-molecule diffusion trajectories were extracted from the .avi file using the ImageJ/Fiji Particle Tracker plugin. Default plugin settings were used, except for kernel radius = 4.0, percentile = 2.5, and link range = 1 (for diffusion constant measurements) or 2 (for counting trajectories). Mathematica scripts (version: 13.2.1.0 for Mac OS X 64 bit) were used to analyze individual videos for their diffusion parameters and average number of total molecules per frame, and to export trajectories for incorporation into an Excel (version 16.101.3) table containing all trajectories measured for a given diffuser. Excel was used to calculate mean step length timecourses, to extract the step lengths used to construct step length frequency distributions, and to calculate frequencies of very long steps. Finally, Mathematica scripts were used to (i) plot frequency distributions of single trajectory diffusion constants, (ii) construct step length frequency distributions and carry out single and multi-Rayleigh distribution analysis to determine diffusion constants, and (iii) plot trajectories.

**Figure 3.**
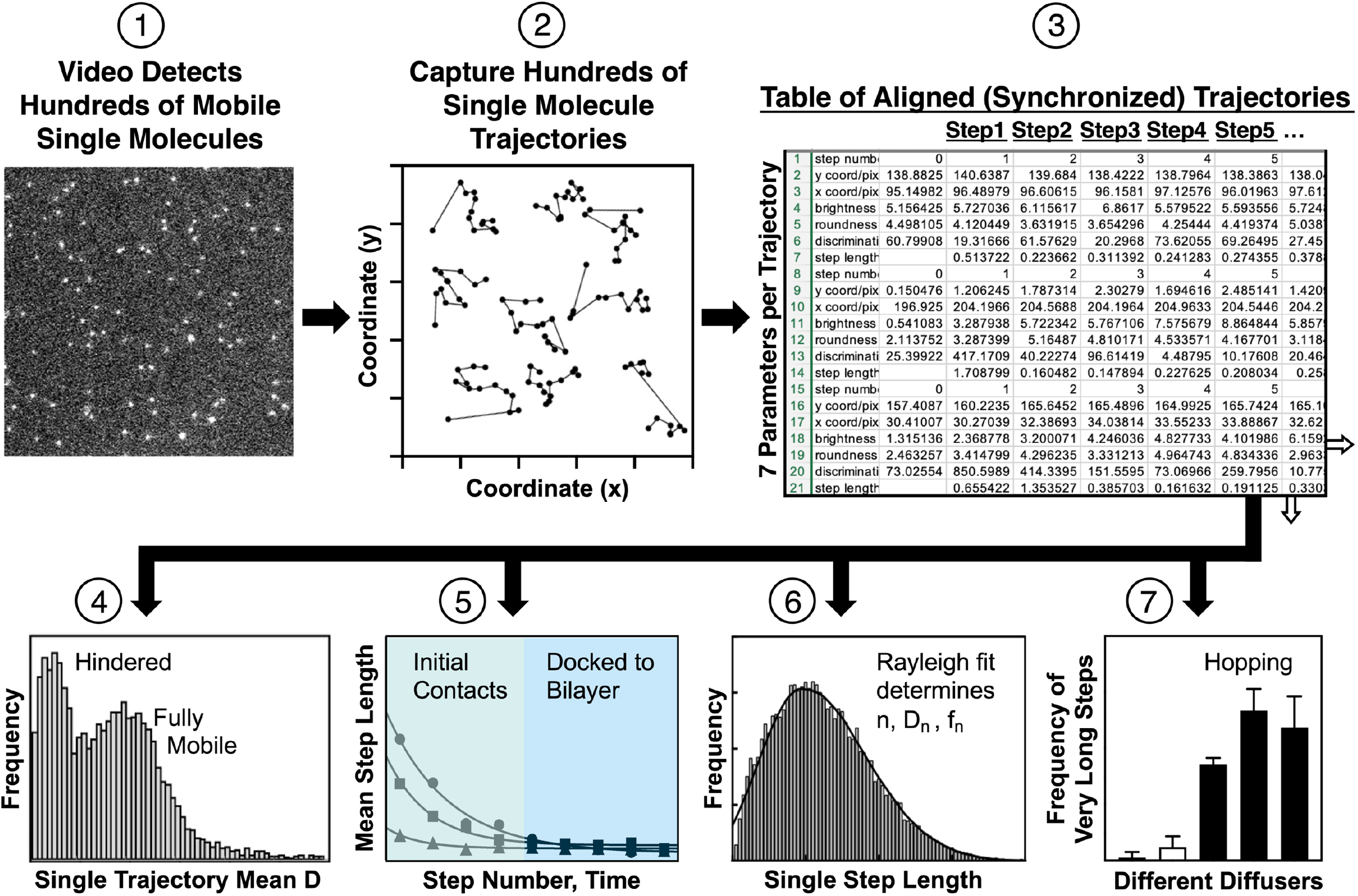
Schematic work flow of single molecule data analysis. **(1)** Single molecule TIRFM is employed to acquire 5 sec video imaging a population of fluorescent diffusers moving on a supported lipid bilayer. **(2)** The video contains hundreds of single molecule trajectories that are acquired and tabulated using particle tracking software (ImageJ/Fiji and Particle Tracker). **(3)** A Mathematica script filters out minor populations of spurious trajectories arising from immobilized or contaminating fluors, then exports the remaining trajectories for assembly into a master Excel summary file containing the filtered trajectories from all videos of a given diffuser, now virtually synchronized by aligning their first steps in column 1. For given trajectory, the table includes its stepwise total brightness, the coordinates of its brightness centroid, the displacement (step length) of this centroid from that of the previous step, and other parameters. **(4)** A Mathematica script imports the master table, calculates the average diffusion constant (Mean D) of each trajectory, and plots the resulting pool of single trajectory diffusion constants as a frequency distribution that resolves hindered from mobile diffusers. **(5)** An Excel script calculates the mean step length for each column or step in the master table, excluding column 1 since the first step is generally not full length. The mean step length is plotted vs time (t = step number x 20 msec) to yield a step length timecourse. For membrane docking proteins this timecourse resolves into two phases corresponding to initial bilayer contacts yielding a decaying mean step length, followed by stable bilayer docking phase exhibiting a relatively constant mean step length. **(6)** For a selected range of steps in the master table, a Mathematica script inputs the individual step lengths of all trajectories and uses it to build a step length frequency distribution, which is best fit with one or more Rayleigh function(s) to ascertain whether diffusion is homogeneous or heterogeneous, and determine the diffusion constants and relative proportions of subpopulations. **(7)** The desired pool of 20 msec step lengths is tested for the presence of fast, long-range hops by an Excel script that measures the frequency of very long steps, defined by a step length expected to occur at a very low frequency (1/1000) in the Rayleigh distribution of a diffusing lipid exhibiting D ∼ 2.0 µm^2^/s.

The frequency of very long steps relative to that expected for a mobile lipid was calculated as follows. The diffusion constant of mobile lipid was observed to be D = 2.0 µm^2^/s during steps 2-3 and during steps 4-12 (Table 1). The Rayleigh distribution for this D indicates that 0.1% (1/1000) of all step lengths are > 1.1 µm, which was defined as the threshold for very long steps (VLS). For a given diffuser, the Excel “countif” command was used to determine the number of VLS in steps 2-3, and separately in steps 4-12, which was divided by the total number of steps to determine the frequency of VLS. For each diffuser, the measured total frequency of VLS includes the frequency of contaminating VLS due to particle tracking errors, which could be determined directly for the mobile lipid by subtracting its expected Rayleigh VLS frequency (0.1%) from its total VLS frequency. The remaining frequency of VLS due to particle tracking errors was subtracted from the total VLS frequency for each diffuser, yielding its observed VLS frequency, which was then divided by the expected Rayleigh VLS frequency or mobile lipid (0.1%) to yield its observed / expected VLS frequency ratio normalized to mobile lipid. The resulting parameter ranged from 1.0 for the lipid diffuser to 52.8 for a membrane docking protein that exhibits hopping behavior.

**Table 1.** Diffusion constants, relative populations, and very long step frequencies for each diffuser.

| **Parameter\*** | **(DO)PE-LRB** | **(DO)PE-Rab1A** | **PX Domain** | **Autophagy Lipid Kinase** | **Kinase (–) Substrate PI** |
|---|---|---|---|---|---|
| Number of Videos | 38 | 29 | 66 | 32 | 34 |
| Number of Trajectories | 7619 | 4207 | 8798 | 3368 | 4314 |
| D1 (steps 2–3) ± SEM (μm²/s) | 2.030 ± 0.050 | 2.097 ± 0.055 | 1.089 ± 0.036 | 0.718 ± 0.012 | 0.765 ± 0.040 |
| D2 (steps 2–3) ± SEM (μm²/s) | 0.532 ± 0.123 | 0.433 ± 0.021 | 2.320 ± 0.290 | 2.390 ± 0.160 | 2.570 ± 0.280 |
| P1 (steps 2–3) ± SEM (%) | 82.868 ± 3.223 | 72.032 ± 1.240 | 66.548 ± 3.961 | 66.992 ± 6.285 | 68.547 ± 3.595 |
| P2 (steps 2–3) ± SEM (%) | 17.131 ± 3.224 | 27.967 ± 1.241 | 33.451 ± 3.962 | 33.007 ± 6.285 | 31.452 ± 3.595 |
| Obs/Exp VLS Frequency (steps 2–3) ± SEM | 1.000 ± 1.300 | 6.500 ± 4.600 | 30.200 ± 2.700 | 52.800 ± 9.200 | 44.500 ± 13.200 |
| D1 (steps 4–12) ± SEM (μm²/s) | 2.046 ± 0.016 | 2.029 ± 0.041 | 1.330 ± 0.090 | 1.338 ± 0.207 | 0.797 ± 0.065 |
| D2 (steps 4–12) ± SEM (μm²/s) | 0.597 ± 0.063 | 0.582 ± 0.035 | 0.000 ± 0.090 | 0.528 ± 0.127 | 2.144 ± 0.427 |
| P1 (steps 4–12) ± SEM (%) | 82.779 ± 1.503 | 70.118 ± 2.286 | 100 ± 0.100 | 64.162 ± 14.443 | 69.115 ± 9.090 |
| P2 (steps 4–12) ± SEM (%) | 17.221 ± 1.503 | 29.881 ± 2.286 | 0.000 ± 0.100 | 35.837 ± 14.443 | 30.884 ± 9.090 |
| Obs/Exp VLS Frequency (steps 4–12) ± SEM | 1.000 ± 1.300 | 2.900 ± 1.300 | 0.200 ± 3.600 | 4.000 ± 1.700 | 10.900 ± 1.700 |

\* D1 and D2 are the diffusion constants obtained by 2-Rayleigh fit of the step length frequency distribution for steps 2-3 (Figure 7) or steps 4-12 (Figure 8).  P1 and P2 are the relative population sizes of the corresponding diffusion states.  For steps 4-12 of PX domain, a 1-Rayleigh fit was adequate thus D2 and P2 are set to zero with the same uncertainties as D1 and P1, respectively.  The observed frequency / expected frequency of very long steps (> 1.1 μm) is normalized to that expected for a mobile lipid under the present conditions, as calculated in Methods.  Statistical and error analysis were carried out as described in Methods and Figure Legends.

### Reproducibility and Statistics

For each diffuser, 1-2 investigators carried out independent experiments on multiple days to collect 29-66 videos, each 5 sec in length and containing hundreds of single molecule trajectories. Altogether, a total data set of 3000 - 9000 filtered trajectories was obtained for each diffuser. The pooled trajectories were analyzed as outlined in Figure 3, with additional details and statistics provided in the figure legends. Where indicated, levels of significant differences were determined by 2-tailed t-test, or by 1-tailed t-test when testing a difference with clear direction. The exceptions were F-tests carried out to compare 1-vs 2-Rayleigh fits of step length frequency distributions (see Supplemental Figures S2, S3 for details).

## RESULTS

### TIRFM measurements of single molecule diffusion on a target membrane - approach

#### Fluorophores and sensors employed

Total internal reflection fluorescence microscopy (TIRFM) was employed to detect and track single fluorescent molecules diffusing in 2 dimensions on a supported lipid bilayer. Fluorophores possessing high quantum yield and high photostability were used to detect the 4 diffusers (**Figure 1A** and Methods). For (DO)PE lipid and PX domain, fluor was coupled chemically to the headgroup or to a labeling Cys near the N-terminus, respectively. For the docking proteins, a 13-residue ALFA tag (SRLEEELRRRLTE) was first added to the N-terminus of Rab1A or to the N-terminus of the Beclin1 subunit in autophagy lipid kinase. Subsequently, when needed for visualization, an anti-ALFA nanobody possessing chemically coupled fluor was added to generate fluorescent, ALFA-tagged Rab1A or autophagy lipid kinase. The latter on-demand approach to fluorescent tagging (Gotzke et al 2019) significantly increases the flexibility of experimental design.

#### Supported lipid bilayers

Supported lipid bilayers were deposited on the surface of an ultra-clean glass slide and, except where noted, contained the native lipid headgroups essential for PX or autophagy lipid kinase activity (Ohashi et al 2020). All lipids employed possessed the same pair of acyl chains (1,2-dioleoyl-sn-glycero-3-phospholipids (DO), which are common in biological membranes and yield stable supported bilayers characterized by homogeneous, easily quantified lipid diffusion (Ziemba et al 2013; Knight & Falke 2009). The standard lipid mixture was (DO)PC / (DO)PS / (DO)/PE / (DO)PI / (DO)PE-MCC in mole ratios 50:25:18:5:2, respectively that yielded a 30% mole fraction of negatively charged lipids, similar to the native negative charge densities of the autophagophore and phagosome membranes. Studies of the PX domain also included a small mole fraction (< 1%) of its target lipid (DO)PI(3)P where indicated.

#### TIRFM detection of single molecule diffusion trajectories

TIRFM generates an evanescent field of excitation photons up to ∼100 nm off the glass surface yielding localized, membrane-proximal excitation zone (Kudalkar et al 2016). Diffusers in the aqueous phase are moving too rapidly to be imaged by the TIRFM system, while diffusers interacting with the supported bilayer are slowed by orders of magnitude and can be imaged. Thus, the TIRFM method selectively detects and monitors the 2D movements of fluorescent diffusers associated, at least transiently, with the supported lipid bilayer. A given 5-second video typically contains hundreds of single molecule trajectories that are captured by single molecule tracking, and subsequently analyzed using a suite of software applications (**Figure 3**).

#### Initial analysis of diffusion trajectories

**Figure 4A** displays representative 2D diffusion trajectories for each of the 4 diffusers on the standard lipid mixture, as well as those for autophagy lipid kinase on a modified lipid mixture lacking its substrate lipid (DO)PI. For each of the 5 diffuser-bilayer systems, thousands of trajectories were obtained in dozens of videos obtained on multiple observation days. Analysis of the resulting pooled trajectories began with a calculation estimating the average diffusion constant of each individual trajectory. The resulting single trajectory diffusion constants were used to construct a frequency distribution as displayed in **Figure 4B** for each of the 5 systems. Each diffuser exhibits a dominant population of mobile molecules; in addition, the lipid-anchored diffusers (DO)PE-LRB and (DO)PE-Rab1A each exhibit a smaller subpopulation of hindered molecules. For the fluorescent lipid, the slow diffusion state has been shown to arise from hindered lipid diffusion in the leaflet against the glass (Vasquez et al 2014). For the lipid-anchored G-protein Rab1A, we observed the appearance of aggregates that grew with time during video collection, yielding a significant subpopulation of molecules with hindered diffusion within the aggregates. Previous studies of lipid-anchored Ras have demonstrated photochemical formation of G-protein oligomers in TIRFM studies (Lin et al 2014). In the present study only bilayers with anchored Rab1A and their controls underwent an overnight coupling reaction that may have yielded membrane defects that recruit G-proteins. Alternatively, some studies suggest that G-proteins can natively oligomerize (Sianoya et al 2025). Thus the observed Rab1A aggregates may be artifactual or potentially interesting. The present study focused solely on the larger, mobile populations of the (DO)PE-LRB and (DO)PE-Rab1A diffusers, and ignored the smaller, hindered populations. In addition, standard filters were employed to eliminate trajectories due to damaged, completely immobilized diffusers and fluorescent contaminants (see Methods).

**Figure 4.**
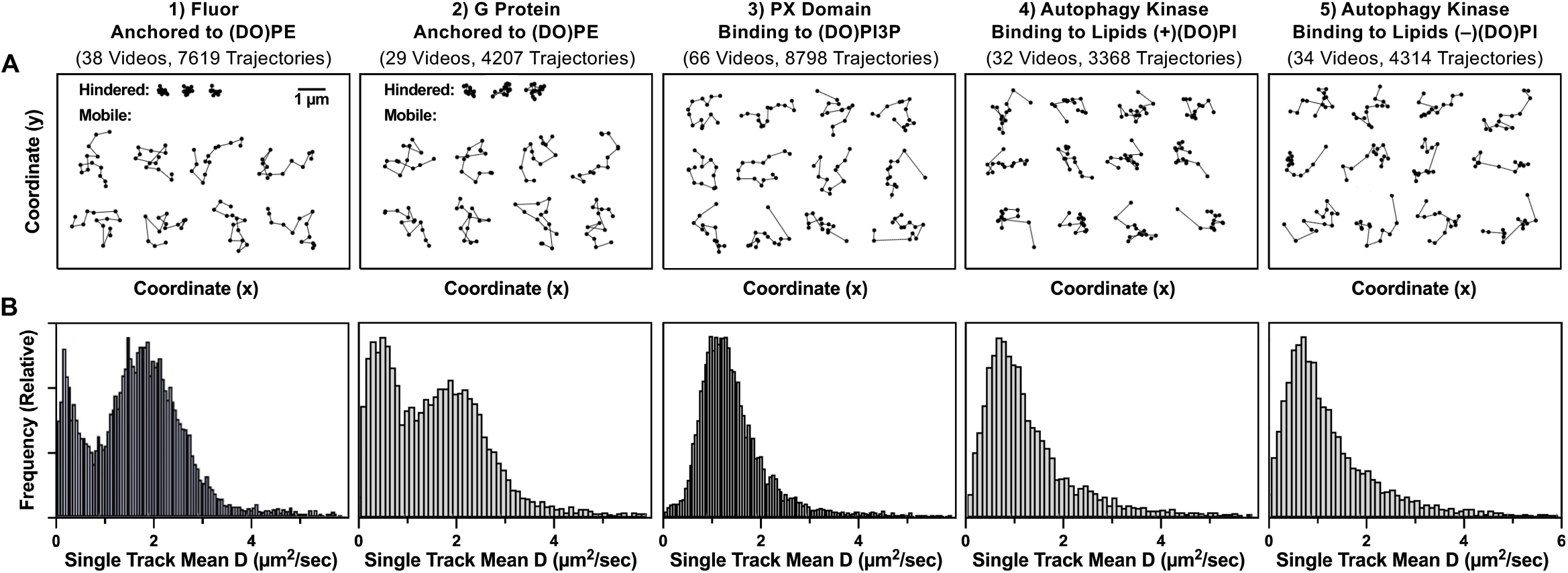
Representative single molecule trajectories, and the distribution of single trajectory diffusion constants, for each diffuser. **A)** Shown are representative 14-step trajectories for each diffuser, and the total number of videos and filtered trajectories collected. Each segment is the displacement between consecutive 20 msec frames. Significant populations of hindered trajectories were observed for both fluor and G-protein anchored to (DO)PE lipid. The supported bilayer was (DO)PC / (DO)PS / (DO)PE / (DO)PI / (DO)PE-MCC, mole percent 50:25:18:5:2. For autophagy lipid kinase, two sets of data were collected on bilayers containing and lacking (DO)PI, respectively. The buffer was 25 mM HEPES, 75 mM KCl, 75 mM NaCl, 2.0 mM Mg^2+^, 75 µM Mn^2+^, 5 mM glutathione, 10 µg/mL BSA, 1 mM Mg-ATP, pH 7.4, and the temperature was T = 21.4° C ± 0.5 °C. B) The distribution of single trajectory (track) diffusion constants for each diffuser and condition, illustrating the low-D hindered populations for the two lipid anchored diffusers. All other populations are mobile.

#### Measurement of equilibrium membrane densities, and use in testing PX domain and autophagy lipid kinase for native function

A generalizable single molecule TIRFM method was developed to quantify the membrane density of each diffuser over a wide density range, thereby facilitating quantitative activity and specificity studies. For the lipid anchored fluor (DO)PE-LRB, for which the labeling efficiency is ∼100%, the average density of (DO)PE-LRB molecules is equal to the density of detected fluors. Thus the fluor density must be adjusted carefully to remain within the range of countable densities. In contrast, fluorescent sensor proteins such as PX domain and anti-ALFA nanobody can be employed over a wide range of target densities by adjusting their concentration, and thus their fractional occupation of the target. Herein the PX domain and anti-ALFA nanobody were employed at sub-saturating concentrations to ensure that the density of membrane-bound, fluorescent sensor-target complexes remained in a countable range – saturating sensor would have yielded fluorescent sensor-target densities too high to count, as well as a high fluorescent background that would interfere with imaging. The observed, sub-saturating sensor-target density is easily converted to a the total target density using the known K_D_ value for each sensor binding to its target lipid or protein (K_D_ = 120 ± 20 nM for p40Phox PX domain binding to PI(3)P (Stahelin et al 2003); K_D_ = 5 nM for anti-ALFA nanobody binding to ALFA tag (data not shown)), from which the fraction of target occupied by sensor was calculated and transformed to the total target density.

This single molecule approach was used to measure the membrane-bound densities of PX domain and autophagy lipid kinase on bilayers of different lipid compositions, in order to test their lipid specificity and functionality (**Figure 5**). In Figure 5A the PX domain displays its native high affinity binding to its target lipid (DO)PI(3)P, such that little or no membrane-bound PX domain is observed in the absence of the target lipid. The equilibrium density of PX domain-(DO)PI(3)P complex is significantly increased by the addition of 25 mole percent background anionic lipid ((DO)PS) from (2.16 ± 0.06) x 10^4 to (2.71 ± 0.04) x 10^4 molecules per frame (n = 46, 42 respectively; p < 0.0001). Similarly, the average bound state lifetime of the PX domain-(DO)PI(3)P complex is subtly but significantly increased by the addition of anionic (DO)PS from 0.69 ± 0.05 to 0.74 ± 0.07 sec (n = 46, 42 respectively; p < 0.0002), reflecting only a small decrease in the PX domain off-rate. The observation that a 29% increase in PX domain binding density is associated with only a 7% slower off-rate indicates that most of the density increase arises from a faster PX domain on-rate for binding to (DO)PI(3)P target lipid as the negative charge of the bilayer is increased. These findings support the hypothesis that an electrostatic search mechanism enhances the on-rate for PX domain binding to (DO)PI(3)P, and we predict the fold enhancement will increase at lower (DO)PI(3)P densities.

**Figure 5.**
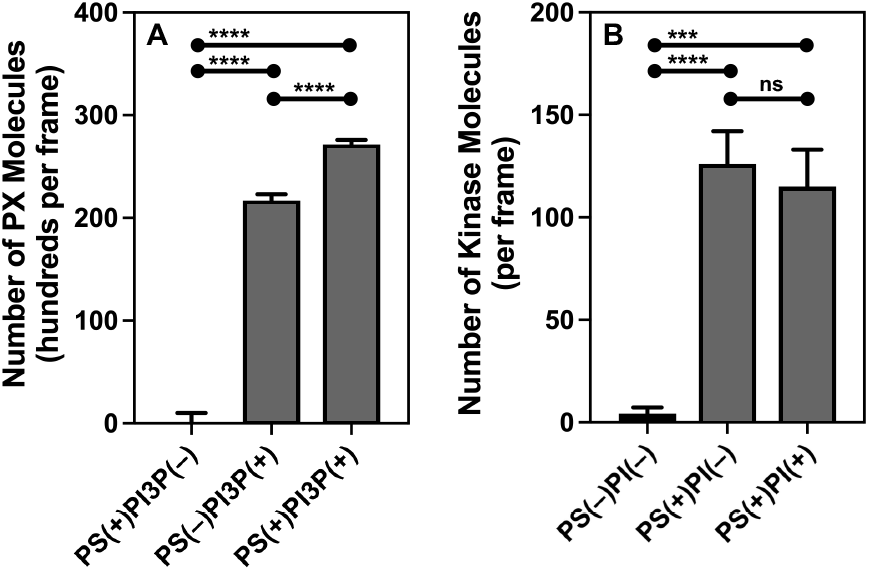
Single molecule, membrane-bound density measurements for PX domain and autophagy lipid kinase illustrate their lipid dependence. The membrane bound density of each indicated diffuser was measured by analysis of single molecule TIRFM data (see Methods). **A)** Shown is the average number of membrane-bound PX domains per frame as measured by single molecule TIRFM. The PX domain requires its target lipid for binding, supplied here as (DO)PI(3)P, while the addition of the anionic lipid (DO)PS further significantly increases binding. **B)** The average number of membrane-bound autophagy lipid kinase molecules per frame, measured by single molecule TIRFM. Autophagy lipid kinase requires anionic lipid for binding, supplied here as (DO)PS. Further addition of its substrate lipid (DO)PI has little or no effect on binding. Only a small fraction of the lipid kinase population is bound to the bilayer, thus the minimal effect of (DO)PI addition is not due to exhaustion of the free lipid kinase pool. The supported bilayer was (DO)PC / (DO)PS / (DO)PE / (DO)PI / (DO)PE-MCC, mole percent 50:25:18:5:2; where (DO)PC was used to replace the (DO)PS and/or (DO)PI lipids that were removed. Other conditions as in Figure 4. Statistics: each bar plots the average number of molecules per frame ± S.E.M. for a given diffuser and condition, calculated as follows. A Mathematica script was used to determine the mean density of molecules per frame for each of n videos. These n means were then averaged to give the global mean ± S.E.M., where n = 37-46 for PX domain and n = 5-7 for kinase.

**Figure 5B** displays smTIRFM measurements of autophagy lipid kinase membrane-bound densities on bilayers of different lipid compositions. The results indicate that the addition of 25 mole percent background anionic lipid ((DO)PS) to increase the bilayer negative charge yields a highly significant, ≥ 28-fold increase in the density of membrane-bound autophagy lipid kinase (p < 0.0001). In fact, the density of bound kinase is negligible before the addition of the anionic lipid. In contrast, in the presence of the 25 mole percent background anionic lipid, the further addition of 5 mole percent anionic substrate lipid (DO)PI has no significant effect on the density of membrane-bound autophagy lipid kinase. Such low affinity for substrate is consistent with the predictions of a recent model for regulation of autophagy lipid kinase catalytic activity (Cook et al, 2025). It follows that autophagy lipid kinase possesses an intrinsic bilayer-docking mechanism that does not require the presence of substrate lipid PI. Despite its low PI affinity, we know that the membrane-bound kinase can bind (DO)PI since slow but significant lipid kinase activity was detected during the smTIRFM experiments (which included ATP (Supplemental Fig. S1)).

### PX domain and autophagy lipid kinase display long hops during their initial contacts with a target bilayer

#### Mean step length trajectories reveal longer average step lengths for membrane docking proteins making their initial bilayer contacts

For each diffuser, a mean step length timecourse was calculated to investigate how average 20 msec step lengths change with time after the molecule first contacts the bilayer. The method begins by generating a table of step lengths in which each row m lists the lengths of consecutive steps for the mth single molecule trajectory in the list, while each column j lists the length of the jth step for each trajectory in the list. The trajectories are aligned with the length of step 1 in column 1, thereby generating virtual synchronization of the entire set of trajectories. Column 1 is omitted from subsequent analyses because step 1 usually is not a full step. This organization of the data enables simple analysis of the properties of individual steps, or groups of steps.

**Figure 6** plots the mean lengths of consecutive 20 msec steps 2-12 against time. The relatively constant mean step length timecourses of the lipid-anchored diffusers (DO)PE-LRB and (DO)PE-Rab1A)) are qualitatively simpler than the two-phase timecourses exhibited by the docking proteins (PX domain and autophagy lipid kinases). The mean step lengths of lipid-anchored diffusers are relatively constant with time, consistent with their permanent coupling to lipid that ensures a consistent mode of diffusion throughout the timecourse. In contrast, the docking proteins exhibit longer mean step lengths during early steps 2-3, before decaying to a roughly constant mean step length. The resulting two-phase timecourses suggest that early steps during the initial membrane may include longer hops until stable bilayer association is achieved, at which point the average steps become shorter and relatively constant.

**Figure 6.**
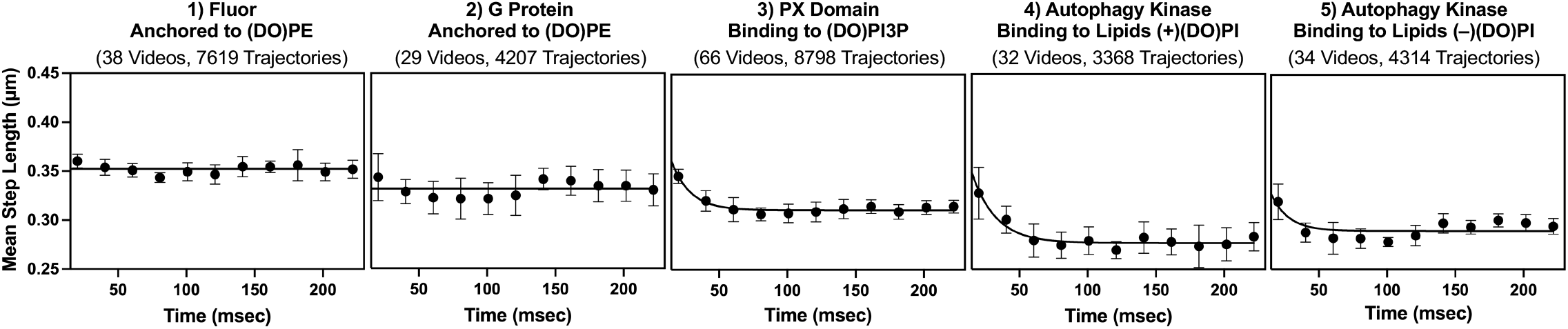
Mean step length timecourses reveal biphasic kinetics for the membrane association of docking proteins, but not for lipid-anchored diffusers. Filtered trajectories were aligned with their first frame at time zero, yielding synchronization of their timecourses. The average step length was determined for each step and plotted (in each case step 1 was omitted since it is generally not a full step). The resulting mean step length timecourses reveal that, for the two lipid-anchored diffusers, the mean step length is relatively constant over the ∼200 msec observation period. In contrast, the timecourses of the membrane docking proteins are biphasic, with the initial steps within 50 msec of membrane contact exhibiting longer mean step lengths, followed by stable bilayer association and relatively constant, asymptotic mean step lengths. The asymptotic mean step lengths are a measure of bulk average mobility for each set of trajectories. The fluorescent lipid exhibits a longer asymptotic mean step length than the docking proteins, while the shortest asymptotic mean step length is that of autophagy lipid kinase in the presence of its substrate lipid (DO)PI. These bulk mean mobilities are in good agreement with the diffusion constants measured by Rayleigh analysis (see below). Experimental conditions as in Figure 4. Statistics: data points are mean step lengths ± S.D. for n = 9, where standard deviations were determined by dividing the full set of trajectories into 9 blocks, then determining the mean step length for each block, and averaging the 9 block means to determine the block S.D. (This block S.D. more accurately defines the level of uncertainty in the mean step length than the raw S.D. determined for the full set of trajectories, which is very large due to the high level of stochasticity in diffusion step lengths).

#### Rayleigh analysis of early steps 2-3 reveals two diffusion states, including a significant subpopulation of possible hops for docking proteins making initial bilayer contacts

For each diffuser, the lengths of steps 2 and 3 were extracted from the step length table to generate a step length frequency distribution. Each distribution, in turn, is subjected to Rayleigh analysis to learn more about the diffusion states present in these early steps. For a population of homogeneous diffusers with a single diffusion constant (D), the step length frequency distribution can be fit with a single Rayleigh function corresponding to that D value. If n different diffusion states are present with different D values, then a sum of n distinct Rayleigh functions is needed to fit the distribution (Schutz et al 1997).

**Figure 7A** displays the step length frequency distribution for each diffuser as well as the best-fit Rayleigh curve; in each case a sum of two Rayleigh functions is needed to adequately fit the distribution. **Figure 7B,C** and **Table 1** compare the D values and fractional populations of the two diffusion states detected for each diffuser. During steps 2-3, both of the lipid-anchored diffusers (DO)PE-LRB and (DO)PE-Rab1A exhibit a major population of fast steps (D ∼ 2 µm^2^/s) and a minor population (16 - 28 % of total steps) of slow steps associated with hindered diffusion (Figures 3,6; Table 1). For both of these lipid-anchored diffusers, herein we focus on the mobile populations that exhibit the diffusion parameters expected for a single (DO)PE lipid. The diffusion constant of a single lipid in the mobile leaflet of the supported lipid bilayer is defined simply by its frictional drag against the bilayer (Knight et al 2010; Ziemba et al 2012; Ziemba & Falke 2013; Camley et al 2015). Since the diffusion constants of mobile (DO)PE-LRB and (DO)PE-Rab1A molecules are the same, within error, the G-protein Rab1A possesses no detected protein frictional drag against the bilayer and thus is adequately described as a soluble protein tethered to the lipid.

**Figure 7.**
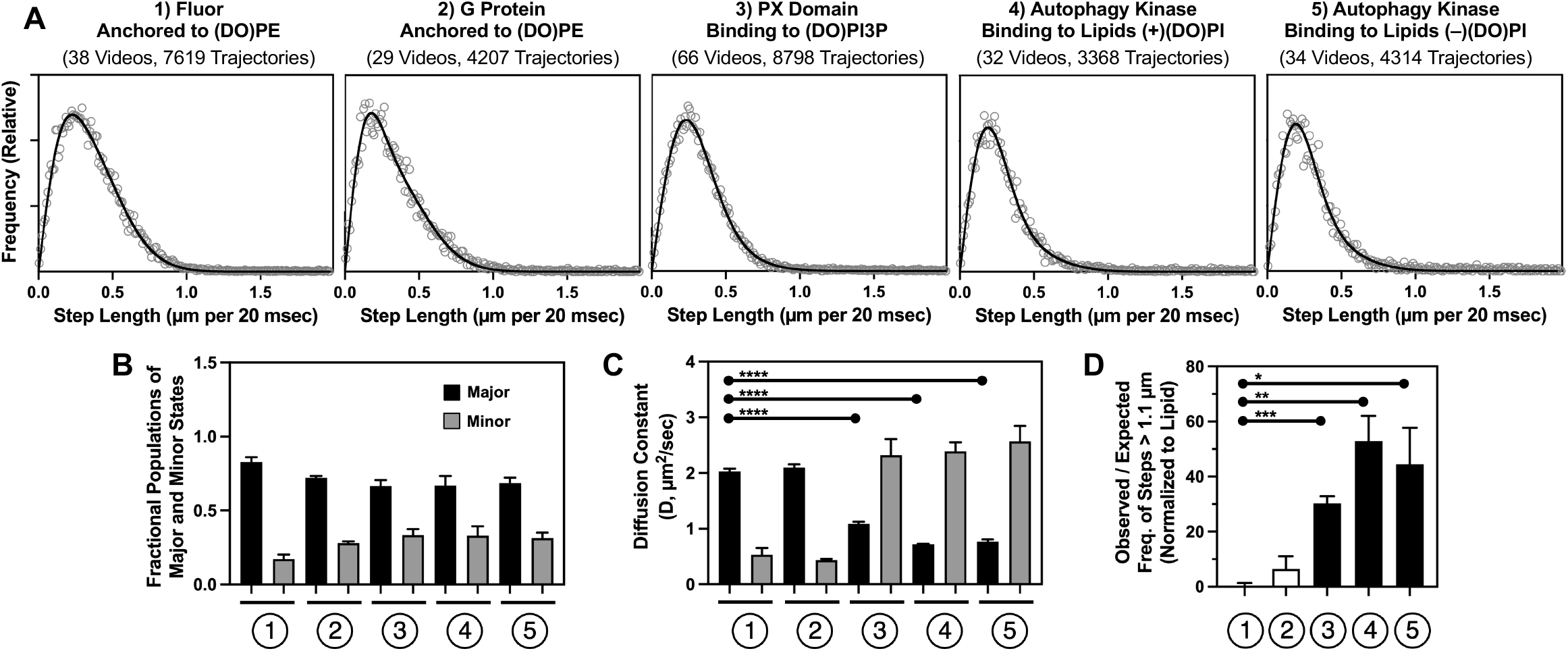
Diffuser motional components during early steps 2-3: Rayleigh analysis of diffusion constants, and very long step frequency analysis of hopping events. **A)** Step lengths were extracted from steps 2-3 of the filtered trajectories for the indicated diffuser and used to construct a step length frequency distribution. The distribution was best fit with 1 or 2 Rayleigh functions and in each case an adequate fit required 2 Rayleigh functions indicating the presence of 2 diffusion components, as justified by F-test analysis (Supplementary Figure S2). **B)** and **C)** Each 2 Rayleigh best fit yielded the fractional populations and diffusion constants of the two diffusion components. In each case, a major population (black bar) and minor population (grey bar) is observed. For the lipid-anchored diffusers the minor, slow diffusing components possess well understood hindered diffusion. For each docking protein, a major population significantly slower than mobile lipid is observed, as well as a minor population as fast or perhaps faster than mobile lipid. **D)** Very long step analysis yielded the observed / expected frequencies of very long steps (> 1.1 µm), normalized to the ratio observed for the mobile lipid (f = 1.0). The frequency of very long steps for lipid-anchored Rab1 was not significantly different from lipid, while the docking proteins exhibited very long step frequencies 30-50 fold higher than lipid, indicating the presence of a hopping component in each case. Experimental conditions as in Figure 4. Statistics: (B,C) Each bar plots the indicated mean value ± S.E.M. for n = 3, where the standard error of the mean was determined by dividing the full set of step lengths for steps 2-3 into 3 blocks, then determining the mean step length for each block, then averaging the 3 block means to determine the block S.D. and S.E.M. for n = 3. (D) Each bar plots the mean ± S.E.M. of the observed frequency / expected frequency ratio of very long steps for the indicated diffuser, normalized to the ratio determined for mobile lipid in the present studies (see Methods). The standard error of the mean was determined by dividing the full set of step lengths for steps 2-3 into 3 blocks, then determining the normalized observed frequency / expected frequency ratio of very long steps for each block, then averaging the 3 block means to determine the block S.D. and S.E.M. for n = 3.

In contrast, during steps 2-3 both PX domain and autophagy lipid kinase exhibit a major population of slow steps (D ≤ 1.2 µm^2^/s), as well as a significant subpopulation (31 - 34 % of the total steps) of fast steps (D ≥ 2 µm^2^/s) (Figure 7, Table 1). Thus, both docking proteins exhibit 2 diffusion states during the initial bilayer encounter. The slow state diffuses more slowly than lipids, indicating the membrane-docked protein-lipid complex experiences frictional drag against the bilayer arising from its bound lipid and its bilayer-contacting residues (Knight et al 2010; Ziemba et al 2012; Ziemba & Falke 2013; Camley et al 2015). In contrast, the fast state diffuses as fast, or perhaps faster, than the 2 lipid anchored diffusers, suggesting the possibility of hopping events across the bilayer surface that enable faster 2D translation than lipid diffusion within the bilayer. Notably, the presence of (DO)PI has no significant effect on either the slow or fast diffusion state of autophagy lipid kinase, suggesting minimal interaction with its substrate lipid during the initial membrane contacts.

#### Analysis of steps 2-3 reveals a high propensity of very long steps for membrane docking proteins making initial bilayer contacts, thus providing strong evidence for hops

To independently test the hypothesis that the docking proteins exhibit hopping behavior when they first contact the bilayer, we compared the frequencies of very long steps (> 1.1 µm per 20 msec) during steps 2-3 for each diffuser (**Figure 7D**). All frequencies were normalized to the expected frequency of very long steps for the fluorescent lipid (DO)PE-LRB, which for a diffusion constant of D = 2 µm^2^/s is predicted to undergo such a step once every 1000 steps. The resulting normalized very long step frequency for (DO)PE-LRB (f = 1 ± 2) is not significantly different from that of the lipid-anchored G-protein (DO)PE-Rab1 (f = 6 ± 4, p = 0.33). In contrast, the PX domain in the presence of (DO)PI(3)P, as well as the autophagy lipid kinase in the presence and absence of substrate (DO)PI, exhibit significantly greater normalized very long step frequences (f = 26 ± 2, 40 ± 10, and 45 ± 8 respectively, p < 0.001 to p < 0.05) compared to (DO)PE-LRB (f = 1 ± 1.3). These findings provide direct evidence that, during the initial bilayer contact steps 2-3, the PX domain and autophagy lipid kinase possess mode(s) of movement that enable faster translation across the bilayer surface than observed for lipid diffusion in the bilayer. Herein we operationally define translational modes that yield significantly higher frequencies of very long steps than observed for (DO)PE-LRB as “hopping” modes. In hopping mode, the diffuser spends at least part of a given 20 msec step diffusing through aqueous solution between transient, nonspecific electrostatic contacts with the bilayer. The ∼10^2^ fold lower viscosity of the aqueous phase relative to the bilayer enables a ∼10-fold increase in mean square displacement, thereby explaining how one or more electrostatic hops during a given 20 msec frame can yield a step longer than 1.1 µm.

### PX domain and autophagy lipid kinase exhibit increased bilayer friction and limited rebinding events in the membrane-docked state

#### Mean step length analysis of steps 4-12 reveals bulk characteristics of the membrane-docked state

After the initial steps 2-3, the mean step length timecourses of all the diffusers are relatively constant, but significant differences are observed between their asymptotic amplitudes. The lipid-anchored diffusers exhibit larger asymptotic mean step length values than the membrane docking proteins PX and autophagy lipid kinase. It follows that after steps 2-3, the docking proteins establish significant bilayer contacts that, on average, yield increased frictional drag against the bilayer relative to a simple lipid diffuser. The smallest mean step length amplitude is observed for autophagy lipid kinase in the presence of its substrate lipid (DO)PI, implying a detectable protein-substrate interaction that further increases frictional drag against the bilayer. Note, however, that the asymptotic mean step length values are bulk averages that can hide multiple diffusion states.

#### Rayleigh analysis of membrane-docked steps 4-12 reveals multiple diffusion states, including evidence for sensitivity to substrate lipid

For each diffuser, all step lengths for steps 4 through 12 were extracted from the step length table to generate a step length frequency distribution, which was subjected to Rayleigh analysis. **Figure 8A** displays the resulting step length frequency distribution and best-fit Rayleigh curve for each diffuser. For the PX domain a single Rayleigh function was adequate to fit the distribution, while for the other 4 systems two Rayleigh functions were needed to adequately fit the distribution as indicated by F-test. **Figure 8B,C** and **Table 1** compare the D values and fractional populations of the two diffusion states detected for each diffuser.

**Figure 8.**
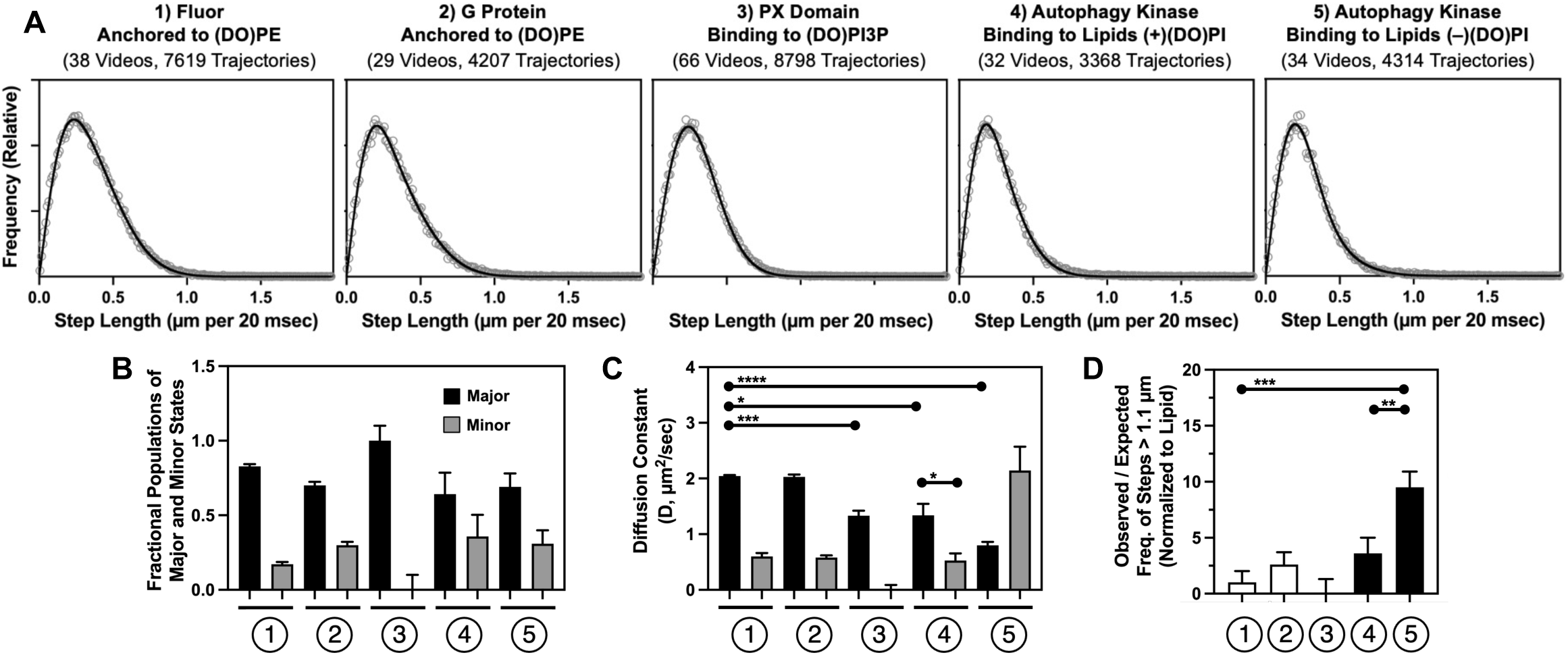
Diffuser motional components during steps 4-12: Rayleigh analysis of diffusion constants, and very long step frequency analysis of hopping events. **A)** Step lengths were extracted from steps 4-12 of the filtered trajectories for the indicated diffuser and used to construct a step length frequency distribution. The distribution was best fit with 1 or 2 Rayleigh functions. For PX domain a single Rayleigh provided adequate fit, while all others required a 2 Rayleigh fit indicating they possess 2 diffusion components (Supplementary Figure S3). **B)** and **C)** Each Rayleigh best fit yielded the indicated fractional populations and diffusion constants, where the black bar indicates the major population. For the lipid-anchored diffusers the minor, slow diffusing components possess well understood hindered diffusion. For each docking protein, the major population continues, as in steps 2-3, to exhibit significantly slower diffusion than mobile lipid. For autophagy lipid kinase, the diffusion of the minor component is significantly slower in the presence of substrate lipid (DO)PI, while in the absence of substrate lipid the minor component is as fast or faster as the mobile lipid population. **D)** Very long step analysis yielded the frequencies of very long steps, normalized to the frequency observed for the mobile lipid (f = 1.0). Only the autophagy lipid kinase in the absence of substrate lipid exhibited a significantly higher frequency of VLS than mobile lipid, indicating the presence of a hopping component in this case. Experimental conditions as in Figure 4. Statistics: (B,C) Each bar plots the indicated mean value ± S.E.M. for n = 3, where the standard error of the mean was determined by dividing the full set of step lengths for steps 4-12 into 3 blocks, then determining the mean step length for each block, then averaging the 3 block means to determine the block S.D. and S.E.M. for n = 3. (D) For the indicated diffuser, each bar plots the mean ± S.E.M (n = 9) of the observed frequency / expected frequency ratio of very long steps measured for steps 4-12 for the indicated diffuser, normalized to the mean ratio determined for mobile lipid in steps 4-12 in the present study (see Methods).

During steps 4-12, as observed in steps 2-3, both of the lipid-anchored diffusers (DO)PE-LRB and (DO)PE-Rab1A continue to exhibit a major population of fast steps (D ∼ 2 µm^2^/s) and a minor population (18 - 30 % of total steps) of slow steps arising from hindered diffusion (Figures 3,6; Table 1). As before, the mobile populations of both lipid anchored diffusers retain the expected characteristics of a single lipid diffusing in the bilayer. Since the mobile G-protein exhibits the same diffusion constant, within error, as the mobile fluorescent lipid, it follows there is again no detectable interaction of the G protein with the bilayer other than its covalent tether to the (DO)PE headgroup.

After the initial membrane contacts of steps 2-3, the PX domain switches from two-state diffusion to a single diffusion constant during steps 4-12. The resulting diffusion constant (D = 1.33 ± 0.09 µm^2^/s) is not significantly different than that of the slow state observed in steps 2-3 (D = 1.09 ± 0.04 µm^2^/s). In contrast, the fast diffusion component observed in steps 2-3 becomes negligible in steps 4-12, suggesting that the hopping component has been eliminated or is no longer detectable. The simplest explanation is that the slow diffusion constant observed in both the early and late steps arises from PX domain stably bound to its (DO)PI(3)P target lipid, with approximately 2/3 of the PX domain population achieving this target lipid-docked state in steps 2-3 and virtually the entire population acquiring this state in steps 4-12. The diffusion constant of the target lipid-docked state is significantly slower than that observed for the fluorescent lipid (2.05 ± 0.02 µm^2^/s, p < 0.001), indicating that the PX domain-(DO)PI(3)P complex experiences frictional drag against the bilayer arising from both its bound lipid and substantial protein-bilayer contacts (Knight et al 2010; Ziemba et al 2012; Ziemba & Falke 2013; Camley et al 2015).

In the absence of substrate lipid (DO)PI, during steps 4-12, the autophagy lipid kinase continues to exhibit two diffusion states with parameters similar to those observed in steps 2-3 in the absence of substrate lipid. The major population state of steps 4-12 (D = 0.8 ± 0.1 µm^2^/s) is within error of the predominant state of steps 2-3 (D = 0.76 ± 0.04 µm^2^/s), and in both cases, the major state is the slower component. Similarly, the minor state in steps 4-12 (D = 2.1 ± 0.4 µm^2^/s) is within error of the corresponding state in steps 2-3 (D = 2.6 ± 0.3 µm^2^/s), in both cases representing the fast component. The simplest explanation is that the major slow state observed in both the early and late steps is a bilayer-associated state in which autophagy lipid kinase is docked to the bilayer through docking motif(s) other than the substrate binding site. The minor fast state identified in steps 4-12 may represent a continuation of the fast, long-step hopping mode of steps 2-3 (see above) although further analysis of very long step frequencies is needed to test this possibility (see next section).

In the presence of substrate (DO)PI, the autophagy lipid kinase continues to exhibit 2 diffusion states during steps 4-12. In contrast to the other fast state diffusion constants observed for autophagy lipid kinase, the fast state diffusion constant observed for steps 4-12 in the presence of substrate lipid is unique (1.3 ± 0.2 µm^2^/s) and is the major state. It is significantly faster than the autophagy lipid kinase slow state diffusion constants observed for early steps 2-3 in the absence and presence of substrate (0.8 ± 0.1 and 0.7 ± 0.1 µm^2^/s yielding p < 0.05 and 0.03, respectively). It is significantly slower than the autophagy lipid kinase fast state diffusion constants observed for early steps in the absence and presence of substrate (2.6 ± 0.3 and 2.4 ± 0.2, µm^2^/s yielding p < 0.05 and 0.01, respectively). It is also significantly slower than lipid diffusion (2.04 ± 0.02 µm^2^/s, p < 0.02). The minor slow state observed for autophagy lipid kinase in the presence of substrate (DO)PI may have a decreased diffusion constant (D = 0.5 ± 0.1 µm^2^/s) relative to that observed in early steps 2-3 (D = 0.71 ± 0.01 µm^2^/s), although this effect is just short of significance (p = 0.051). Similarly, the presence of substrate (DO)PI may slow diffusion in steps 4-12 relative to that observed in the absence of (DO)PI (D = 0.8 ± 0.1 µm^2^/s), although this effect is again not quite significant (p = 0.051). All these slow state diffusion constants remain significantly slower than lipid diffusion (D = 2.05 ± 0.02 µm^2^/s, p < 0.001), providing strong evidence that the interaction between autophagy lipid kinase and lipids generates substantial frictional drag against the bilayer in all cases.

#### Analysis of membrane-docked steps 4-12 reveals enhanced propensity of very long steps for membrane docking proteins in specific cases

To independently test for the presence of hopping behavior after the initial bilayer contacts, we compared the frequencies of very long steps (> 1.1 µm per 20 msec) during steps 4-12 for all diffusers, again normalizing all frequencies to that of the fluorescent lipid (DO)PE-LRB (f = 1 ± 2) (**Figure 8**). For the lipid-anchored G-protein the frequency of very long steps (DO)PE-Rab1 (f = 3 ± 2) is not significantly different (p = 0.52) from fluorescent lipid. Similarly, for PX domain (f = 0 ± 4), as well as for autophagy lipid kinase in the presence of substrate (f = 4 ± 2), the frequency of very long steps is not significantly different from (DO)PE-LRB (p = 0.83, 0.35, respectively). In contrast, autophagy lipid kinase in the absence of substrate lipid (11 ± 2) exhibits a significantly higher frequency of very long steps than both (DO)PE-LRB (p < 0.01) and autophagy lipid kinase in the presence of substrate (p < 0.05). These findings provide evidence that after the initial bilayer contacts in steps 2-3, where hopping is universally detected for PX domain and autophagy lipid kinase, the presence of a specific target lipid ((DO)PI(3)P for PX domain or (DO)PI for autophagy lipid kinase) virtually eliminates the hopping mode during steps 4-12, or suppresses its amplitude so it is no longer detectable via very long step analysis. In contrast, when its substrate lipid is absent, the autophagy lipid kinase hopping mode continues to be detected during steps 4-12, albeit at a lower frequency, and is hypothesized to arise from unbinding-hopping-rebinding events.

## DISCUSSION

The present comparison of single molecule diffusion trajectories on a target lipid bilayer reveals major differences between the trajectories of lipid-anchored diffusers and peripheral membrane docking proteins. The findings also reveal novel features of the membrane dynamics of both PX domain and autophagy lipid kinase, with important implications for their functions.

The lipid-anchored diffusers, specifically the fluorescent lipid (DO)PE-LRB and the lipid-anchored G protein (DO)PE-Rab1A, each exhibit a dominant mobile population of single molecules (Figure 4) that display a diffusion constant representative of lipid diffusion in the bilayer (Table 1). Since the mobile population of the lipid-anchored G protein exhibits the same diffusion behavior as the mobile fluorescent lipid, it follows that the G protein and its flexible hypervariable region exhibit no detectable frictional drag against the bilayer, and thus that they lack protein-bilayer contacts.

The membrane docking reactions of the peripheral membrane proteins PX domain and autophagy lipid kinase exhibit sensitivity to the bilayer lipid composition (Figure 5), as well as distinct early and late kinetic phases (Figure 6). PX domain requires its target lipid (DO)PI(3)P for detectable binding and exhibits a significant increase in target lipid affinity when the bilayer contains 25% (DO)PS as an anionic background lipid (Figure 5A). The increased affinity for (DO)PI(3)P triggered by (DO)PS in the bilayer stems predominantly from an increased on-rate for target lipid binding linked to the increased density of negatively charge lipid in the bilayer, consistent with an electrostatic search mechanism that speeds target lipid acquisition. **Figure 2** provides a schematic view of a representative single PX domain trajectory. Analysis of a set of 8798 trajectories reveals that during the initial membrane contacts in steps 2-3, up to 1/3 of the population diffusion steps are longer and faster than allowed by lipid diffusion and exhibit the features expected for electrostatic hopping across the bilayer surface (Figures 6,7, Table 1). The other 2/3 of the steps are shorter and slower than lipid diffusion, indicating these steps generally arise from the stably docked PX-(DO)PI(3)P protein-lipid complex. Within ∼ 50 msec after the initial membrane contact, the fast diffusion state and very long steps are no longer detected and the diffusion behavior of the membrane-associated PX domain population becomes homogeneous, having been fully converted to the stably docked PX-(DO)PI(3)P complex. This complex exhibits slower-than-lipid diffusion in the bilayer owing to its frictional drag against the bilayer generated by both the lipid and by membrane docking loops of the protein that penetrate the bilayer (Figure 1B), as previously described for other peripheral membrane proteins (Knight & Falke 2009, Knight et al 2010, Ziemba & Falke 2013, Ziemba et al 2014, Ziemba et al 2016, Buckles et al 2017, Gordon et al 2023). In the stable, membrane-docked state, the lack of fast diffusion and very long steps indicates that unbinding-hopping-rebinding events (Figure 2) are rare and/or are not easily detected at the density of (DO)PI(3)P employed.

Autophagy lipid kinase requires anionic background lipid for stable bilayer association (Figure 5B), and exhibits multiple diffusion states. The absolute requirement for a negatively charged bilayer surface, together with the observation of long autophagy lipid kinase displacements that are faster than lipid diffusion, provide strong evidence that the kinase possesses an electrostatic search mechanism like the PX domain. Altogether 3368 trajectories were obtained on bilayers containing the substrate lipid (DO)PI, and 4314 trajectories on bilayers lacking (DO)PI. Analysis reveals that both in the presence and absence of (DO)PI, the first ∼50 msec of contacts include, like the PX domain, both a fast state accounting for approximately 1/3 of the steps and a slower state accounting for the remaining 2/3 of the steps (Figures 6,7, Table 1). The observation that the fast and slow diffusion states are indistinguishable on bilayers containing and lacking substrate lipid (DO)PI indicates that none of these early diffusion states involve binding to (DO)PI. Given that autophagy lipid kinase possesses two positively charged membrane docking feet, the fast state could involve one or more modes of translation across the bilayer surface including one-footed hops and two-footed skating. The slow diffusion state arises from frictional drag against the bilayer, which controls the diffusion constant of bilayer-associated proteins, with frictional contributions from a combination of bound lipids and protein-bilayer contacts. Since PI binding is not involved, the non-catalytic foot is implicated. Based on previous studies on the relationship between D and frictional drag against the bilayer (Knight et al 2010; Ziemba et al 2012; Ziemba & Falke 2013; Camley et al 2015), the diffusion constant of the slow diffusion state is equivalent to the frictional drag of up to 3 lipid molecules, but more likely arises from 1-2 bound lipids plus protein-bilayer contacts that generate additional drag. (DO)PS is the likely candidate for bound lipid since it is the major anionic lipid in the bilayer and its removal largely eliminates autophagy lipid kinase binding to the bilayer. Positively charged and/or hydrophobic residues of the kinase are hypothesized to provide the substantial, additional protein frictional drag required to explain the observed slow diffusion constant. Thus, the simplest model for the slow diffusion state proposes that the non-catalytic foot dominates the membrane contacts with up to 2 bound (DO)PS molecules, the ALPS helix, and the hydrophobic finger (Fan et al 2011, Ohashi et al 2020) all contributing frictional drag.

In the stable, bilayer-docked state the autophagy lipid kinase exhibits altered translational behaviors, especially in the presence of its substrate lipid (DO)PI (Figures 6,8, Table 1). In the presence of substrate (DO)PI, a new fast state representing the majority of the population is observed with a diffusion constant significantly slower than lipid diffusion, but significantly faster than the slow diffusion states exhibited by this protein. The new fast state diffusion constant is similar to that of the one-footed PX domain-(DO)PI(3)P protein-lipid complex, and no long displacements due to autophagy lipid kinase hopping/skating are detected. Thus the the simplest hypothesis proposes the new state arises from a one-footed state with the catalytic foot docked to its substrate lipid (DO)PI, yielding a frictional drag and diffusion constant similar to the PX domain-(DO)PI(3)P complex. Besides the bound (DO)PI lipid, additional frictional drag is likely provided by basic and aromatic residues of the catalytic foot that contact the bilayer, especially the C-terminal helix of the VPS34 subunit (Brier et al 2019; Stjepanovic et al 2017). (Again, even for the large kinase, the frictional drag against the aqueous phase is negligible relative to that against the bilayer and can be neglected). In addition to the new fast state, a new slow state appears to diffuse more slowly than the other autophagy lipid kinase slow states, although this slowing is borderline significant (p = 0.051; Figure 8, Table 1). The simplest explanation proposes that the slow state includes contributions from the two-footed state bridging the one-footed states (Figure 9A), which combines the additive frictional drags of both one-footed state. In the absence of substrate lipid, the major slow and minor fast diffusion states observed during the stable bilayer-docked state appear to continue with diffusion constants that are unchanged, within error, from those observed during the early bilayer contacts (Figures 7,8, Table 1). The fast diffusion state is proposed to arise from unbinding-hopping/skating-rebinding events, yielding hopping/skating behavior similar to that detected during initial membrane contacts but with a reduced frequency of very long steps. The slow diffusion state is proposed to be the same one-footed state, with the non-catalytic foot docked to the bilayer, implicated by its diffusion constant as the initial slow state during all early bilayer contacts, both with and without substrate (DO)PI.

**Figure 9.**
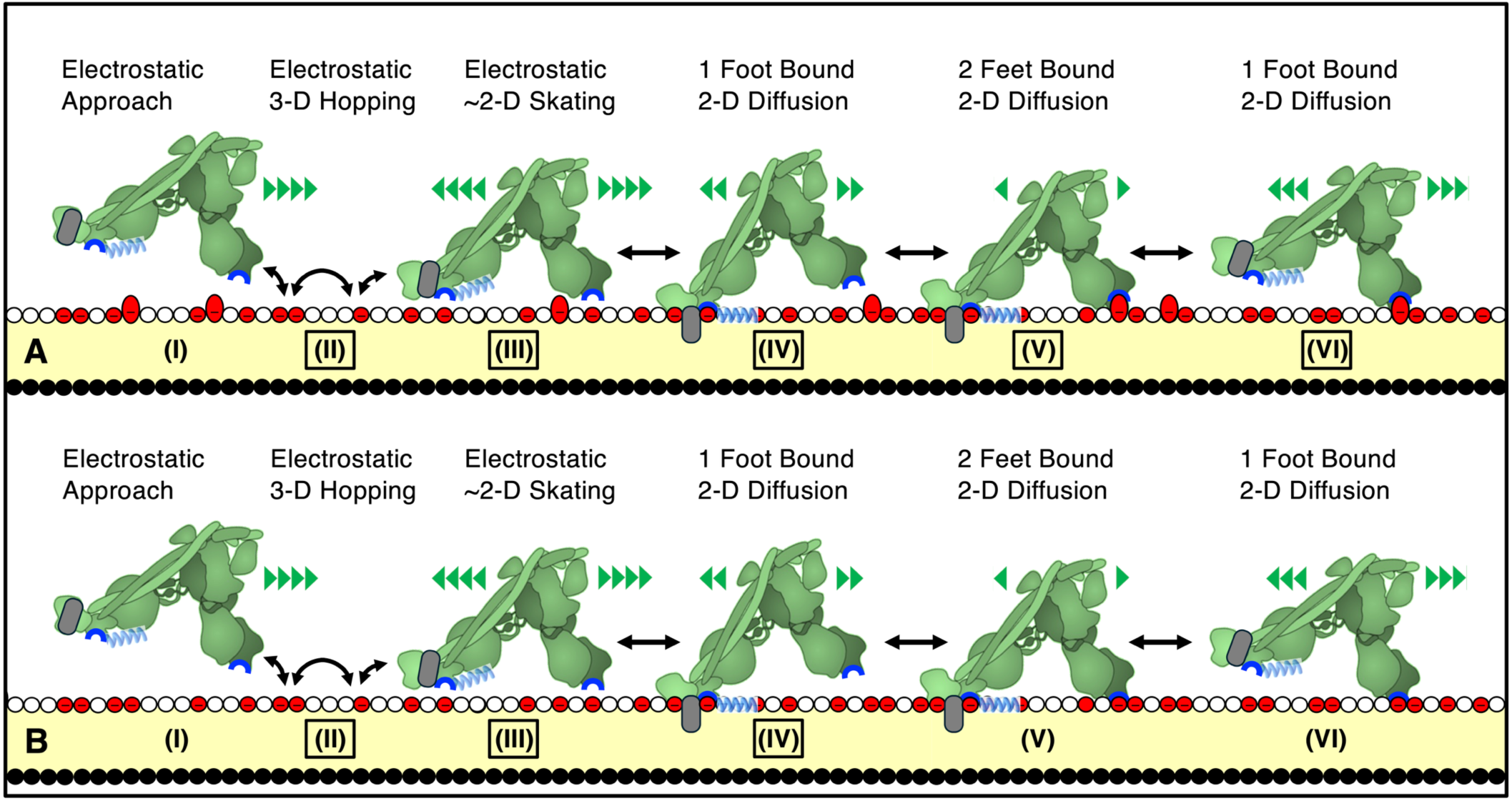
Working model for autophagy lipid kinase docking to the bilayer. Shown are hypothesized movements of the schematic, two-footed autophagy lipid kinase across an anionic bilayer containing phosphatidylserine (PS, red circle headgroups) with (A) or without (B) the kinase substrate phosphatidylinositol (PI, red oval headgroups). The non-catalytic foot (left) possesses the ALPS motif (light blue helix, C-terminus of subunit ATG14L), an aromatic finger (grey cylinder, Beclin subunit), and multiple positive charges (dark blue, supplied in part by ALPS). The catalytic foot (right) possesses the multiple positive charges including the substrate binding site (dark blue, C-terminus of VPS34) as well as several isolated aromatic residues (not shown). States proposed for the observed diffusion constants are boxed. **(States A.II-III, B.II-III; D > 2 µm^2^/s)** For ∼50 msec after the initial bilayer contact, electrostatic interactions enable interconverting one-footed hopping and/or two-footed skating movements across the anionic bilayer surface, with no required order of appearance. These translational movements are faster than lipid diffusion and yield an electrostatic search of the bilayer surface for protein docking targets. For comparison, the diffusion constant of lipid is D ∼ 2 µm^2^/s. **(States A.IV, B.IV; D ∼ 0.8 µm^2^/s)** Within the first ∼50 msec after contact, the electrostatic search ends when the non-catalytic foot docks to the bilayer via ALPS, the aromatic finger, and 1-2 bound PS molecules. In this bilayer-docked state, the 2-dimensional diffusion is slower than lipids owing to the frictional drag of multiple protein elements and bound lipid(s) against the bilayer. **(State A.V; D ∼ 0.5 µm^2^/s)** Despite its low intrinsic affinity for substrate PI, the bilayer-docked protein experiences a high local PI concentration, enabling the catalytic foot to bind a PI headgroup, yielding a two-footed state with additional frictional drag. **(State A.VI; D ∼ 1.3 µm^2^/s)** The two-footed binding mode can transition to a one-footed state in which the catalytic foot is bound to substrate lipid PI, which becomes the predominant state (see Scheme 1A for further discussion). The diffusion constant increases due to the loss of the extensive frictional drag of the non-catalytic foot. **(State B.IV, D ∼0.8 µm^2^/s)** In the absence of substrate lipid, within ∼50 msec of membrane contact the predominant state becomes the one-footed state with the non-catalytic foot docked to the bilayer (B.IV). **(State B.III-II, D > 2 µm^2^/s)** Also observed during this period, but only in the absence of substrate lipid, are fast, very long displacements indicating that one-footed state B.IV undergoes a detectable frequency of unbinding-hopping/skating-rebinding events. (While the schematic models show autophagy lipid kinase “standing” on membrane with the molecular plane normal to the bilayer (Ma et al 2017), this angle likely fluctuates.)

**Figures 9A,B** summarize the resulting working model for the movement of autophagy lipid kinase across a bilayer that either (A) contains or (B) lacks its substrate lipid PI. The model is based on the evidence in Figures 6, 7, 8 and Table 1. The data suggests that the initial bilayer contacts are the same in the presence or absence of substrate PI, indicating that binding to substrate lipid is not involved in these early steps (A.I-IV, B.I-IV). First, the kinase protein executes electrostatic hops, thereby moving faster than lipid diffusion across the bilayer surface (A.II-III, B.II-III). These hops may well include one-footed electrostatic hops and/or two-footed electrostatic skating making use of the net positive charges of both feet. During the first ∼50 msec following the initial membrane contact, the hopping/skating mode is terminated when the non-catalytic foot establishes stable bilayer contacts likely to include PS binding, as well as bilayer penetration of the ALPS helix and aromatic finger. The resulting frictional drag against the bilayer yields a slow diffusion state that translates slower than lipids in the bilayer. By the end of the first ∼50 msec nearly all of the autophagy lipid kinase molecules have achieved this one-footed, bilayer docked state (A.IV, B.IV). Subsequently, autophagy lipid kinase exhibits different behaviors on bilayers containing or lacking substrate lipid PI (Figures 9A and B, respectively). In the presence of substrate PI, the new, faster bilayer-docked diffusion state is proposed to be the one-footed state with the catalytic foot bound to substrate PI (A.VI), which arises via step-wise transitions from the one-footed docking state anchored by the non-catalytic foot (A.IV), to a two-footed docking state in which the catalytic foot binds to substrate lipid (A.V), followed by release of the non-catalytic foot to generate the one-footed state with the catalytic foot bound to its substrate (A.VI). This working model is consistent with both the present observations and with published evidence indicating that autophagy lipid kinase possesses low affinity for substrate in the absence of activating G-protein (Cook et al 2025). In this scenario, the docking of the non-catalytic foot to bilayer facilitates substrate lipid binding to the catalytic foot by greatly increasing the effective local concentration of substrate headgroups. Once formed, the two-footed docking state possesses additional frictional drag (A.V), thereby further slowing the slow state diffusion from D = 0.8 to D = 0.5 µm^2^/s (Figure 8, Table 1), although this reduction is borderline significant (p = 0.051 for one-tailed t-test). The latter slow diffusion constant likely includes contributions from both slow states (A.IV-V) due to their interconversion. The two-footed state is proposed to be conformationally strained, such that it rapidly decays to either one-footed state. When the non-catalytic foot releases, the resulting one-footed state is docked to the bilayer via the PI-bound catalytic foot (A.VI) and exhibits a relatively fast, diffusion constant due to the loss of the extensive frictional drag of the non-catalytic foot (A.IV). No hopping/skating events are detected in this state, suggesting it is a terminal state leading to dissociation from membrane. In the absence of substrate lipid, states involving the membrane-docked catalytic foot (B.V-VI) are not detected. The fast hopping mode detected after the appearance of the bilayer-docked, slow diffusion state is proposed to arise from unbinding-hopping/skating-rebinding events, suggesting this one-footed state can unbind and carry out additional electrostatic searches before rebinding the bilayer (B.IV-II, B.II-IV)).

A puzzle emerges: if the presence of substrate lipid shifts the predominant membrane-docked state from a one-footed state with the non-catalytic foot docked to bilayer (A.IV) to a different one-footed state with the catalytic foot docked to PI on the bilayer (A.VI), one might conclude that the latter one-footed state is thermodynamically favored relative to the other two bilayer-docked states (A.IV, A.V). Yet the density of autophagy lipid kinase on the bilayer is the same, within error, on membranes containing or lacking the substrate lipid (Figure 5), and both feet are protected from aqueous exposure in HDX studies of the bilayer-docked kinase on membranes containing PI (Ohashi et al 2020). We hypothesize the following kinetic scheme that harmonizes these apparently contradictory findings:

**Scheme 1.**
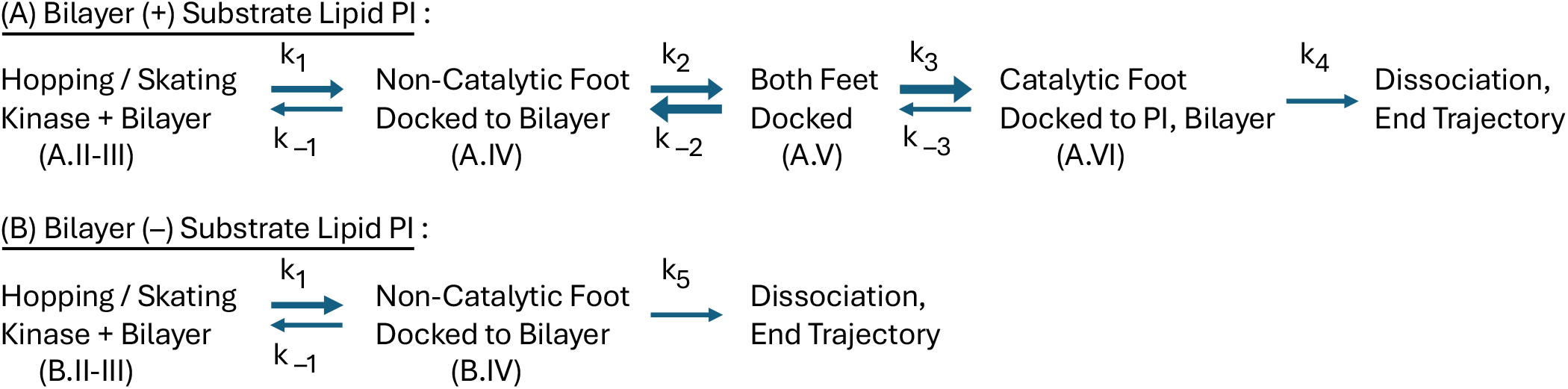

In Scheme 1, the hopping / skating states (A.II-III, B.II-III) interconvert with each other during fast translation across the surface of the bilayer, until the non-catalytic foot docks to the bilayer (A.IV). Scheme 1A proposes that when substrate PI is present at sufficiently high density in the membrane, the initial one-footed docked state (A.IV) is converted to the two-footed docked state (A.V) by the binding of the catalytic foot to PI. The two-footed docked state (A.V) is proposed to be conformationally strained, so it quickly converts either back to the original one-footed docked state (A.IV) or to the other one-footed state with the catalytic foot docked to PI (A.VI). When the forward rate constants k_2_, k_3_ are faster than the reverse rate constant k_–3_ the one-footed docked state bound to PI (A.VI) will be the predominant bilayer-docked state and unbinding-hopping/skating-rebinding events will be rare (Figure 8). Scheme 1B proposes that in the absence of PI, the one-footed state with the non-catalytic foot docked to bilayer is the predominant docked state (B.IV) and can experience unbinding-hopping/skating-rebinding events, as observed (Figure 8). When the rates of dissociation of the two predominant bilayer docked states are similar, or k_4_ ∼ k_5_ for (A.VI) and (B.IV), respectively, then the density of lipid kinase on the bilayer will be similar in the presence and absence of substrate lipid PI, as observed (Figure 5). Moreover, the model can explain the similar partial HDX protection of both feet from solvent in the presence of substrate PI (Ohashi et al 2020), since the diffusion constants indicate that non-catalytic foot residues are more deeply buried in the bilayer, which counterbalances the predicted relative population of states in Scheme 1A (A.V < A.IV < A.VI).

Functionally, the electrostatic translocation modes of PX domain and autophagy lipid kinase are well-suited for speeding encounters with target lipids and proteins on the surfaces of native endosomal membranes. Following phagocytosis, the appearance of target lipid PI(3)P on the phagosome membrane recruits p40Phox via specific binding of its PX domain to the target lipid headgroup. Subsequently, p40Phox binds the membrane-anchored NOX2 NADPH oxidase complex to activate production of reactive oxygen species that neutralize pathogens trapped in the phagosome (Suh et al 2006; Montano-Rendon 2021). The electrostatic hopping-based search mechanism of the p40Phox PX domain is proposed to significantly enhance this front-line defense strategy of innate immunity by increasing the on-rates of p40Phox binding to both its target lipid (PI(3)P) and target protein (NOX2 NADPH oxidase complex), thereby speeding the appearance of a robust ROS burst.

In autophagy, the ability of the kinase to use its two positively charged feet to carry out an electrostatic scan of the membrane surface is proposed to help maximize its on-rate for binding to its membrane-associate activators, including Rab1A, ULK1, GABARAP and NRBF2 (Spoikaite et al 2026; Dessus et al 2026; Chen & Hurley 2025; Mercer et al 2021; Tremel et al 2021; Buckles et al 2020; Young et al 2016; Ohashi et al 2016). Such activators may well be found at low density on the expansive autophagosome membrane, increasing the functional impact of the electrostatic search mechanism. Furthermore, the observed propensity of autophagy lipid kinase to spontaneously form a stable, non-specific bilayer contact in the absence of a high affinity target lipid or protein could, in principle, help distribute the kinase broadly across the autophagosome membrane to ensure widespread distribution of PI(3)P generation centers. The resulting delocalized PI(3)P production is in turn proposed to facilitate the widespread recruitment of PI(3)P-binding regulatory proteins and complexes essential for autophagophore bilayer growth, including WIPI proteins and DFCP1 (Ohashi 2021b).

Further studies are needed to develop detailed kinetic and molecular pictures of the protein-bilayer interactions central to electrostatic scanning by autophagy lipid kinase and the PX domain. The kinetics of switching between the electrostatic scanning and bilayer-docked states may be accessible to two-state kinetic and/or machine-learning analysis of transitions between states in single molecule diffusion tracks. A molecular picture of the nature of protein-membrane contacts stands as a challenging but exciting goal for course-grain simulations of proteins with positively charged feet translating across a negatively charged bilayer surface. Besides furthering a molecular understanding of signaling biology, a biophysical understanding of electrostatic scanning and bilayer docking could facilitate the development of drug delivery systems that more rapidly and efficiently scan cell surfaces to acquire binding targets.

## AUTHOR CONTRIBUTIONS

Conception JJF; experimental design JJF, AMM; biochemistry YO, LL, AMM; TIRFM data collection LL, JJF; development of Mathematica scripts JDK, JJF; development of Excel scripts JJF, LL; data analysis LL, JJF; data interpretation JJF, LL; writing of manuscript JJF, YO,LL; grant support JJF, RLW; helpful conversations JJF, RLW, YO, JDK, LL.

## ACKNOWLEDGEMENTS

The authors gratefully acknowledge funding by NIH Grant R35 GM144346 to JJF, and Cancer Research UK grant DRCPGM\100014 and UKRI Medical Research Council MC_U105184308 grant to RLW. We thank Moshe Gordon for helpful discussions regarding Excel scripts, and Antoine Dessus for the microscale thermophoresis measurement of K_D_ for anti-ALFA nanobody binding to the ALFA tag on G protein and kinase constructs.

---

**Figure S1.**
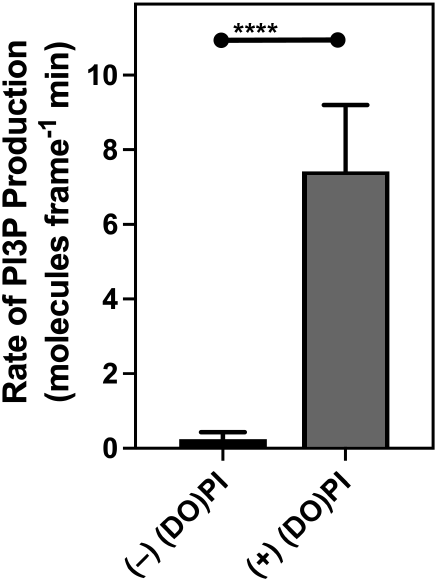
Lipid kinase assay confirms the catalytic activity of the autophagy lipid kinase utilized in the present single molecule TIRFM studies. All single molecule TIRFM measurements were conducted in autophagy lipid kinase activity buffer including ATP, so that whenever substrate lipid (DO)PI was present the kinase was generating product lipid (DO)PI(3)P. The activity was low since the kinase was studied in the absence of its activating G-protein, but kinase activity was measurable over the course of the experiment, which ranged from 110 - 235 min. At the end of the experiment, fluorescent PX domain was added to bind and detect product (DO)PI(3)P molecules, and the total density of product (DO)PI(3)P was calculated from the known PX domain concentration and its K_D_ for binding (DO)PI(3)P. Finally, the total (DO)PI(3)P molecules was divided by the elapsed minutes since adding kinase to generate the plotted rate. The activity in the absence of substrate lipid was at least 60-fold lower, confirming that the kinase retained measurable activity in single molecule TIRFM experiments even in the absence of activating G-protein. The supported bilayer was (DO)PC / (DO)PS / (DO)PE / (DO)PI / (DO)PE-MCC, mole percent 50:25:18:5:2; where (DO)PC was used to replace the (DO)PI lipid removed where indicated. Other experimental conditions as in Figure 4. Statistics: each bar plots the mean rate ± S.D. for n = 42 and n = 55, respectively.

**Figure S2.**
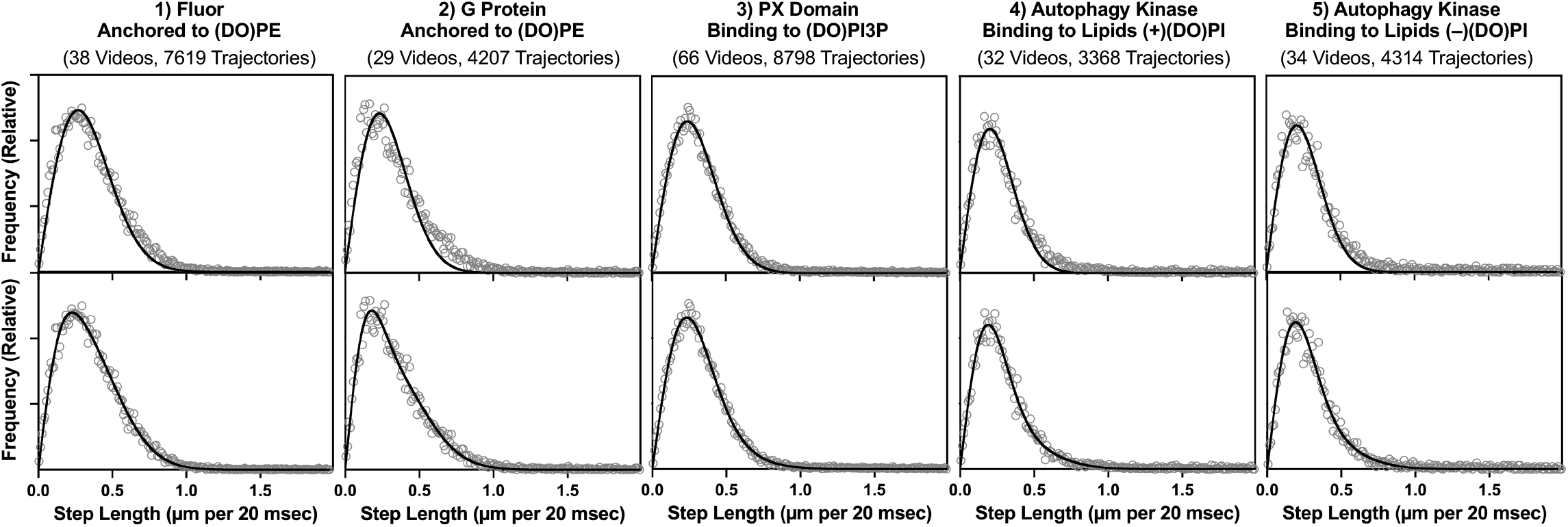
Diffuser motional components during early steps 2-3: Comparison of 1- and 2-Rayleigh best fit analysis. Step lengths were extracted from steps 2-3 of the filtered trajectories for the indicated diffuser and used to construct a step length frequency distribution. The distribution was best fit with 1 (upper) or 2 (lower) Rayleigh functions. In each case an adequate fit required 2 Rayleigh functions indicating the presence of 2 diffusion components, as justified by F-test analysis (p < 0.001).

**Figure S3.**
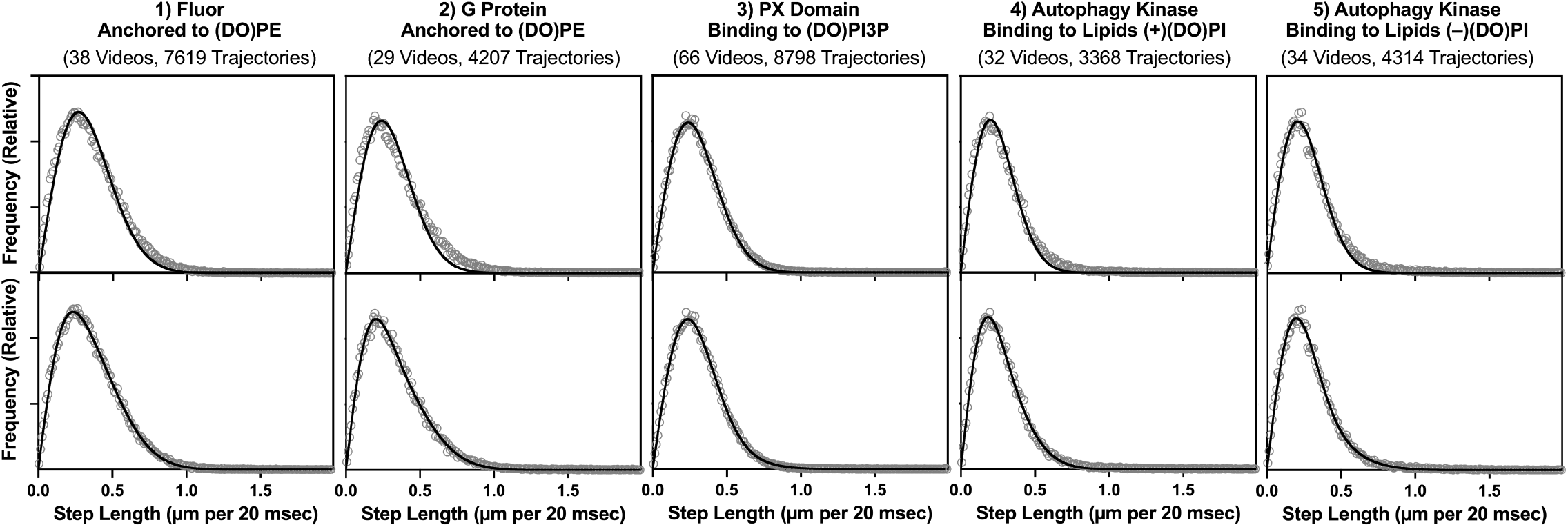
Diffuser motional components during steps 4-12: Comparison of 1- and 2-Rayleigh best fit analysis. Step lengths were extracted from steps 4-12 of the filtered trajectories for the indicated diffuser and used to construct a step length frequency distribution. The distribution was best fit with 1 (upper) or 2 (lower) Rayleigh functions. For PX domain a single Rayleigh provided adequate fit, while all others required a 2 Rayleigh fit indicating they possess 2 diffusion components, as justified by F-test analysis (p < 0.001). In each case, the F value was calculated from the variances (V1, V2) and degrees of freedom (df1, df2) for the 1- and 2-Rayleigh fits of the indicated distributions, respectively.

